# The Data Distillery: A Graph Framework for Semantic Integration and Querying of Biomedical Data

**DOI:** 10.1101/2025.08.11.666099

**Authors:** Taha Mohseni Ahooyi, Benjamin Stear, J. Alan Simmons, Vincent T. Metzger, Praveen Kumar, John Erol Evangelista, Daniel J. B. Clarke, Zhuorui Xie, Heesu Kim, Sherry L. Jenkins, Mano R. Maurya, Srinivasan Ramachandran, Eoin Fahy, Thomas H. Gillespie, Fahim T. Imam, Natallia Kokash, Matthew E. Roth, Robert Fullem, Dubravka Jevtic, Aleks Mihajlovic, Michael Tiemeyer, Clara Bakker, Andrew J. Schroeder, Julia Markowski, Jared Nedzel, Dave D. Hill, James Terry, Christopher Nemarich, Jyl Boline, Peter J. Park, Kristin G. Ardlie, Jeet Vora, Raja Mazumder, Rene Ranzinger, Bernard de Bono, Shankar Subramaniam, Jeffrey S. Grethe, Jeremy J. Yang, Christophe G. Lambert, Adam Resnick, Aleks Milosavljevic, Avi Ma’ayan, Jonathan C. Silverstein, Deanne M. Taylor

## Abstract

The Data Distillery Knowledge Graph (DDKG) is a framework for semantic integration and querying of biomedical data across domains. Built for the NIH Common Fund Data Ecosystem, it supports translational research by linking clinical and experimental datasets in a unified graph model. Clinical standards such as ICD-10, SNOMED, and DrugBank are integrated through UMLS, while genomics and basic science data are structured using ontologies and standards such as HPO, GENCODE, Ensembl, STRING, and ClinVar. The DDKG uses a property graph architecture based on the UBKG infrastructure and supports ontology-based ingestion, identifier normalization, and graph-native querying. The system is modular and can be extended with new datasets or schema modules. We demonstrate its utility for informatics queries across eight use cases, including regulatory variant analysis, tissue-specific expression, biomarker discovery, and cross-species variant prioritization. The DDKG is accessible via a public interface, a programmatic API, and downloadable builds for local use.

## 1. Introduction

Modern biomedical research has produced large multi-modal datasets spanning across genomic variation, gene expression, chromatin structure, metabolomics, protein interactions, and phenotypic traits. These data provide complementary insights into biological systems but are difficult to integrate due to inconsistent formats, annotations, and semantic frameworks. Integrative analyses typically require custom pipelines, manual harmonization, and substantial domain expertise to link across data types—for example, connecting regulatory variants to tissue-specific expression or tracing the metabolic consequences of drug action^1^.

Knowledge graphs (KGs) address these challenges by representing biomedical entities (e.g., genes, diseases, drugs, metabolites) and their relationships (e.g., regulation, interaction, expression) in a structured graph. This supports semantically rich, multi-hop queries that reflect how biologists conceptualize systems: as networks of interrelated components^2–4^. KGs offer a scalable method for integrating disparate datasets using ontologies and controlled vocabularies, enabling complex biological queries across domains in a unified framework ^5–13^

The NIH Common Fund Data Ecosystem (CFDE) aims to improve data interoperability across large-scale initiatives including the 4D Nucleome^14^, exRNA ^15–17^, GlyGen ^18^, GTEx ^19^, HuBMAP^20^, Illuminating the Druggable Genome (IDG) ^21^, Kids First ^22^, LINCS ^23^, Metabolomics Workbench ^24^, MoTrPAC ^25^, and SPARC ^26^. Despite these efforts, data integration across biological types and scales remains challenging because of metadata inconsistencies, identifier conflicts, and heterogeneity in relationship definitions.

The Data Distillery Knowledge Graph (DDKG) was developed to address this gap by integrating over a dozen CFDE datasets into a harmonized, queryable platform for biomedical discovery (**Figure 1**). Built on the UBKG framework and using the Petagraph schema^10^ the DDKG employs more than 180 ontologies and controlled vocabularies to align various types of omics data, including chromatin loops, eQTLs, gene expression, metabolite measurements, drug-target interactions, and disease phenotypes among many others, into a coherent, semantically structured knowledge graph. The DDKG design emphasizes schema extensibility, identifier normalization, and biological interpretability across tissues, conditions, and molecular modalities.

**Figure 1.**
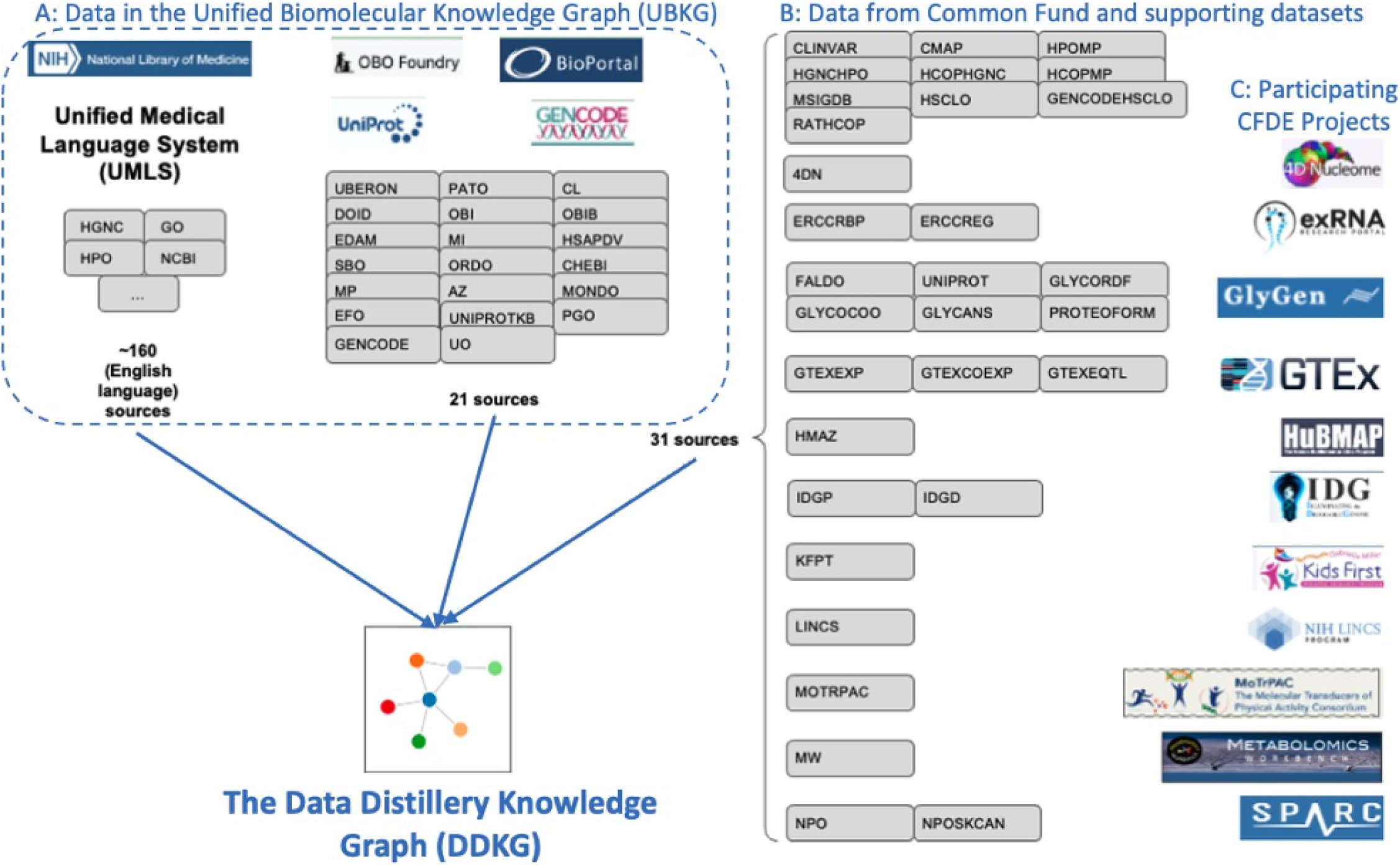
Overview of data sources integrated into the DDKG with the Unified Biomedical Knowledge Graph (UBKG). **(A)** The DDKG harmonizes diverse biomedical knowledge by incorporating over 180 vocabularies and data sources across clinical, genomic, and ontological domains from the UBKG. This includes: (left) the Unified Medical Language System (UMLS) with over 100 English-language terminologies; (center top) ontologies and controlled vocabularies from BioPortal, OBO Foundry, and GENCODE. **(B)** The DDKG extends the UBKG with data from annotation databases and datasets from NIH Common Fund Data Coordination Centers (DCCs), such as GTEx, HuBMAP, LINCS, Kids First, and others. These sources are harmonized through a property graph schema that supports semantic integration and query-driven exploration in the DDKG.

The DDKG also bridges into machine learning by representing node metadata (e.g., labels, synonyms, cross-references) as graph-connected nodes rather than flat attributes. This structure supports flexible subgraph projection and downstream tasks such as node classification or link prediction. The project provides a repository that includes example scripts for preparing ML-ready inputs, thus preserving the graph’s native property graph warehouse and query capabilities ^27^.

The DDKG is accessible through multiple channels. Complete graph builds, including Docker images and component CSV loading files, are available at ubkg-downloads.xconsortia.org. Access requires a free UMLS license key, which can be obtained by registering at https://uts.nlm.nih.gov/uts/signup-login. For interactive exploration, the DDKG-UI (https://dd-kg-ui.cfde.cloud/) provides a web-based interface with guided queries and visualizations, supporting a curated subset of cross-domain search and retrieval tasks.

The underlying platform for the DDKG has previously been described and evaluated ^10^. This paper describes the DDKG’s construction and application through methods for eight use cases including spatial gene regulation, biomarker discovery, drug prioritization, metabolic analysis, and graph-based clustering. These examples illustrate how the DDKG functions not only as a scalable knowledge graph but as a semantically structured integration platform. It enables biologically grounded querying, supports graph-based machine learning workflows, and lowers the barrier to cross-domain exploration by embedding curated biomedical data into a unified, ontology-linked graph environment.

## 2. Results

The Data Distillery Knowledge Graph (DDKG) enables researchers to query and analyze complex, semantically-integrated datasets ascertained from multiple NIH Common Fund Data Coordination Centers (DCCs). Currently, it incorporates data from 4D Nucleome (4DN), exRNA, GlyGen, GTEx, HuBMAP, Illuminating the Druggable Genome (IDG), Kids First, LINCS, Metabolomics Workbench, MoTrPAC, and SPARC along with several other assertion resources, including GENCODE, IMPC, Ensembl, STRING, MSigDB and ClinVar. By supporting harmonized queries across these diverse domains, the DDKG facilitates new opportunities for cross-modal biological discovery. For example, a user might query chromatin loops from 4DN and evaluate their overlap with GTEx eQTLs to identify candidate regulatory interactions that influence gene expression.

While the DDKG supports broad integrative queries, users must consider how to combine datasets in biologically meaningful ways. Integrating data sources without a clear scientific rationale can produce spurious or uninterpretable results. For instance, a DDKG query merging seemingly unconnected phenomena may yield valid outputs, but the biological relevance would be questionable without a guiding hypothesis. The DDKG does not restrict such queries but assumes the user brings appropriate domain knowledge to contextualize the results.

The following use case methods illustrate how DDKG can support biologically grounded exploration across diverse modalities, including chromatin architecture, extracellular RNA, transcriptomics, and chemical perturbation. Each example has been validated both through local execution in Neo4j Desktop and through deployment in the public DDKG-UI interface ^28^. Query templates and app walkthroughs are provided in the Data Distillery User Guide ^28^.

### 2.1 Method 1: Rapid identification of genomic feature neighborhoods: Regulatory Interactions for Chromatin Loops and eQTLs

Resolving spatial proximity among chromosomally localized features is central to many functional genomics and genetics analyses, but typically requires multi-step workflows involving interval comparisons and annotation joins using tools like GenomicRanges or bedtools. Those workflows require several steps when integrating multiple feature types or higher-order relationships. The DDKG enables a flexible, semantically harmonized graph-based approach for integrating any chromosomally mappable feature, such as eQTLs, chromatin loops, expression values, or phenotypes linked by variant associations. This use case illustrates a generalizable strategy using eQTL–loop integration as a minimal example.

We integrated chromatin loop calls from 4D Nucleome (4DN) and liver eQTLs from GTEx v8, modeling both at 1 kbp resolution using the DDKG’s graph-ready Homo sapiens chromosomal location ontology, HSCLO ^29^. A single query (Online Methods, **Query 1**) identified eQTLs within loop segments—upstream anchor, downstream anchor, or interior—linking each variant to its spatial context and associated gene.

The query finds that eQTLs are overrepresented in anchor regions relative to loop interiors, even after normalizing for segment size (**Figure 2C, 2D**). This enrichment aligns with known regulatory roles of anchor points involving CTCF and cohesin binding ^30–33^. The full list is given in **Supplementary File 1**.

**Figure 2:**
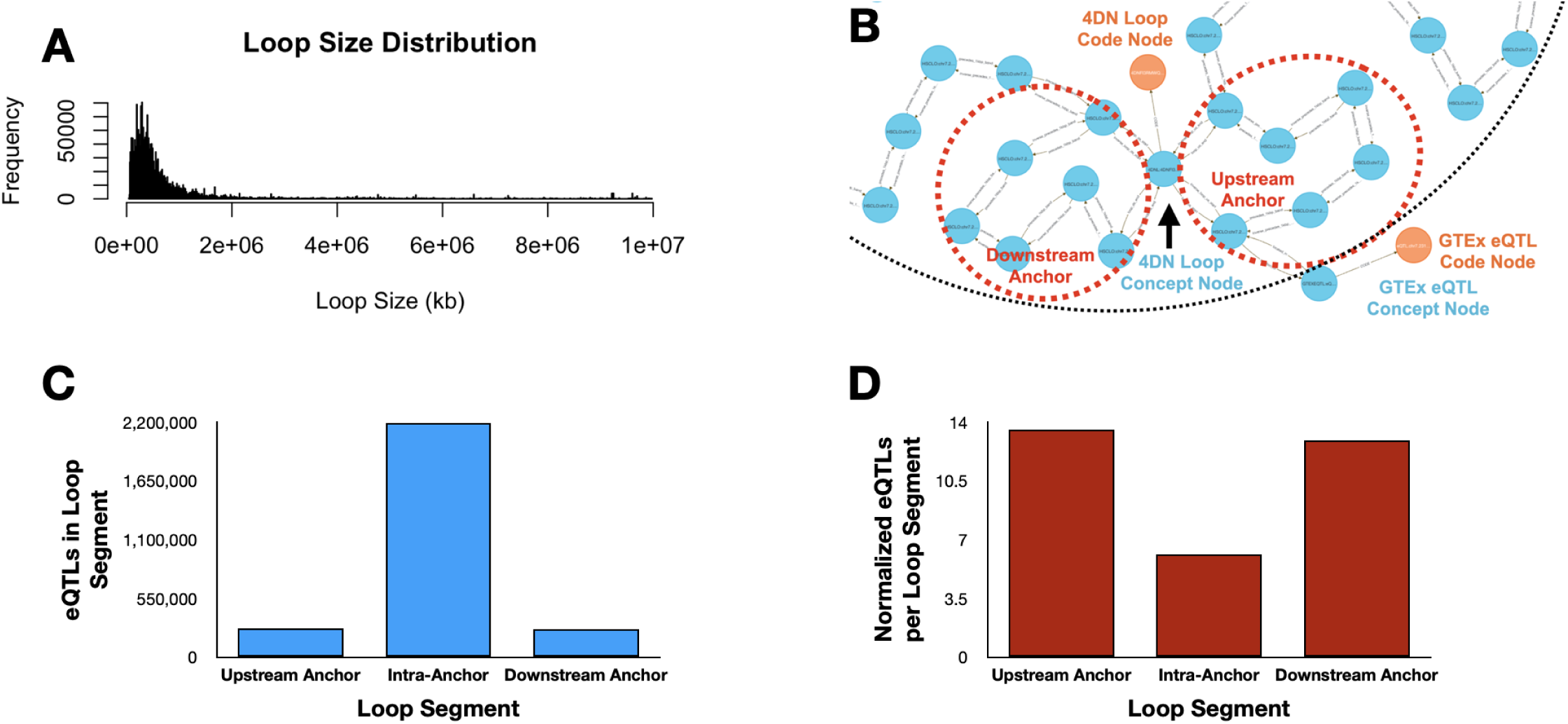
Integration of chromatin loop and eQTL data using the DDKG. A single graph-native query integrates 3D chromatin conformation data from 4DN with GTEx eQTLs, enabling spatial analysis of regulatory variants without external preprocessing. **(A)** Distribution of chromatin loop sizes across 12 4DN datasets. **(B)** Example loop modeled in the DDKG with upstream and downstream anchors (red dashed circles) and an overlapping GTEx eQTL (bottom right). **(C)** Total GTEx eQTLs found in loop segments (upstream, intra-anchor, downstream). **(D)** Normalized frequency of eQTLs per segment, adjusted for segment length, showing enrichment at anchors.

This example demonstrates how the DDKG supports rapid, multi-feature genomic integration within a unified query framework, enabling biologically interpretable analyses that would otherwise require extensive preprocessing and annotation handling.

### 2.2 Method 2: Biomarker Discovery

Identifying biomarkers for non-invasive diagnostics often requires integrating transcriptomic profiles, molecular binding annotations, and disease relevance across multiple data types and biological sources. Traditional workflows for liquid biopsy marker discovery involve merging RNA-seq results with curated databases of RNA-binding proteins (RBPs), filtering for extracellular RNAs (exRNAs) expressed in relevant biofluids, and cross-referencing with disease annotations. These multi-step analyses typically rely on custom scripts and repeated manual reconciliation across tools and formats. The DDKG provides a unified platform for expressing these relationships directly, enabling researchers to identify candidate biomarkers through flexible, multi-modal graph queries. While this use case focuses on disease- and drug-linked exRNAs, the same approach generalizes to any context involving molecular entities traceable through expression, regulation, and localization in biofluids or tissues.

To demonstrate this, we performed two DDKG-based liquid biopsy queries using data from the Extracellular RNA Communication Consortium (exRNA)^15,16^ . In the first example, we searched for exRNAs associated with frontotemporal dementia (HP:0002145) whose loci overlapped known RBP binding sites. The DDKG identified APP, a susceptibility gene with an exRNA locus overlapping a TIAL1 binding site, supporting a candidate non-invasive diagnostic strategy in saliva (see Online Methods, **Figure 7** and **Query 2**).

In **Query 3**, we examined the drug Astemizole, identifying ALCAM as an upregulated gene whose exRNA product overlapped a PTBP1 binding site in cerebrospinal fluid (CSF), a plausible treatment-monitoring biomarker given PTBP1’s high expression in neural tissue (see Online Methods **Figure 8** and **Query 3**).

Both analyses were executed as single queries returning results in seconds, illustrating how graph-native workflows on the DDKG can streamline biomarker discovery across diseases, treatments, and biofluids.

### 2.3 Method 3: GTEx queries for Tissue-Specific Expression

Integrating gene expression data with functional annotations to study tissue-specific patterns is a common task in integrative biology, but it often requires reconciling inconsistent identifiers, tissue ontologies, and annotation formats. Analysts working with GTEx typically extract expression matrices from flat files, align them with curated gene lists, and script custom joins to incorporate other sources of biological context.. The DDKG simplifies this process by semantically linking expression, anatomy, and gene function in a unified graph.

Although this method focuses on joining GTEx data to glycosylation enzymes, the same framework generalizes to joining any source data through gene sets and supports integration with resources like STRING, MSigDB, IDG, LINCS, or user-defined annotations.

To explore glycosylation patterns across tissues, we used **Query 4** (Online Methods) to combine GTEx expression data with enzyme classifications from GlyGen. Glycosylation is essential for protein folding, signaling, and immune recognition, and its dysregulation is implicated in cancer, neurodegeneration, and congenital disorders.

We queried 46 glycoenzyme genes across 54 GTEx tissues and retrieved 2,484 tissue–gene–reaction relationships, modeled as semantically linked Concept nodes. These links were exported and then used to build a log-transformed, hierarchically clustered matrix and visualization using the R package pheatmap^34^ (**Figure 3**).

**Figure 3.**
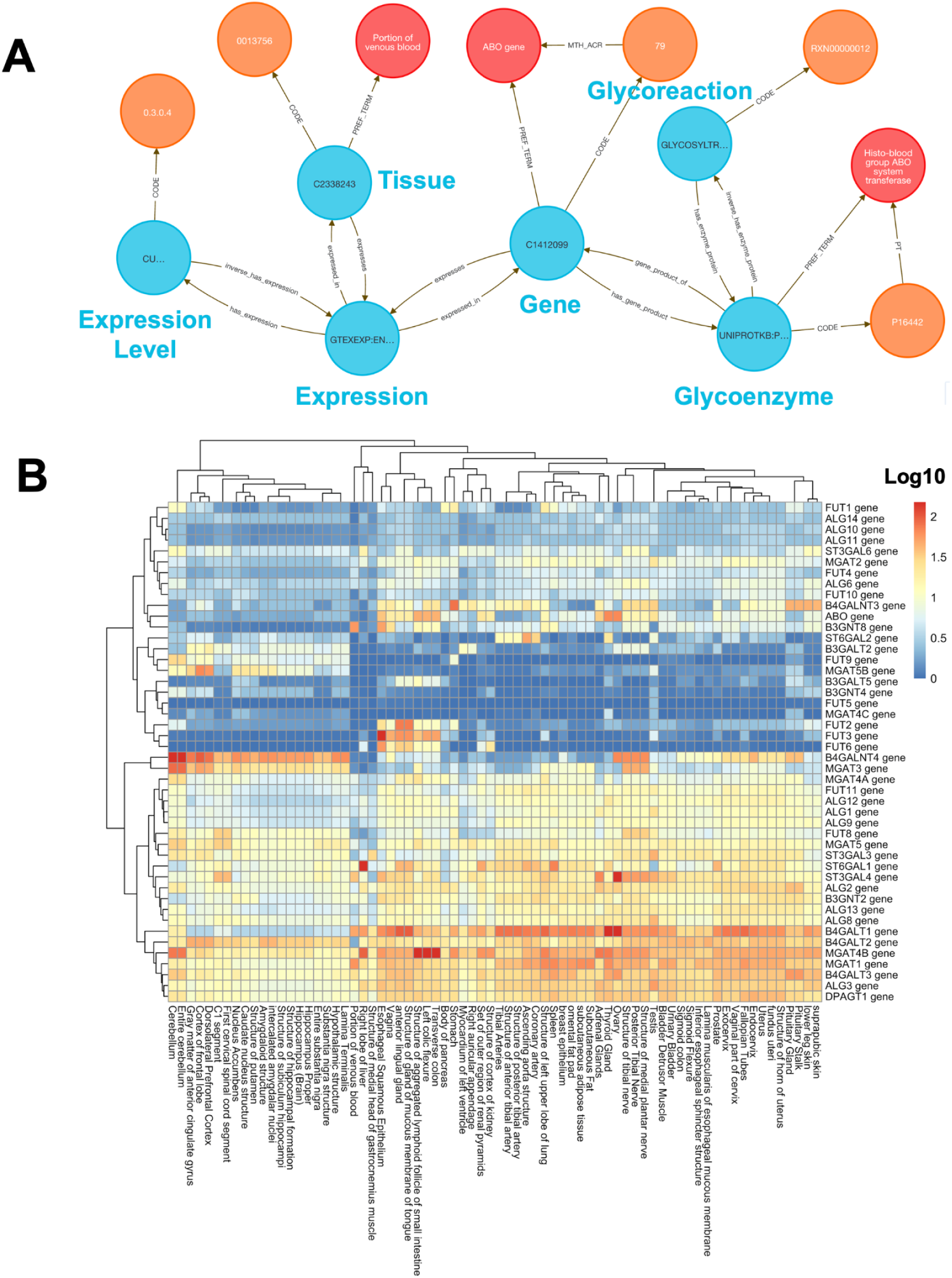
(A) An example graph view showing the connection between glycoenzyme genes and human tissues through the GTEx expressions of the genes encoding glycosylation-associated genes provided by Glygen data (Query 4). (B) Heatmap visualization of tissue-wide expressions (log10 of the mean TPMs) of 46 major glycoenzyme-encoding genes illustrates tissue- and organ-specific variations.

**Figure 4:**
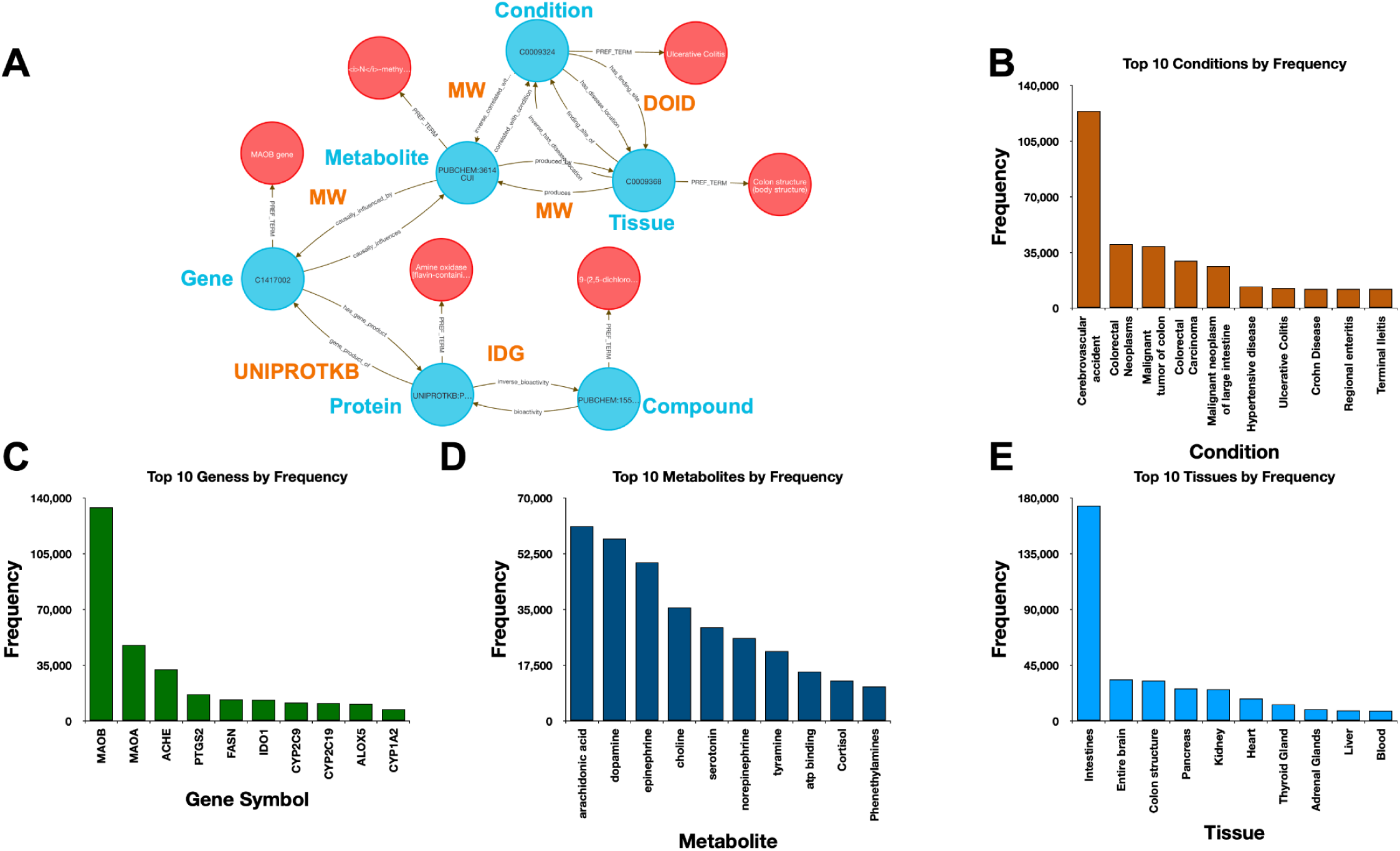
Joint querying assertions defined by Metabolomics Workbench (MW) and IDG. (A) An example graph view of output depicting the cross-walk between a human gene (MAOB), a metabolite, condition (Ulcerative Colitis) and tissue (Colon Structure) and the same gene protein product modulation by a bioactive compound (see Online Methods, Query 8). Top 10 (B) conditions, (C) genes, (D) metabolites and (E) conditions based on frequency in 450,000 output instances as a result of the query.

**Figure 5:**
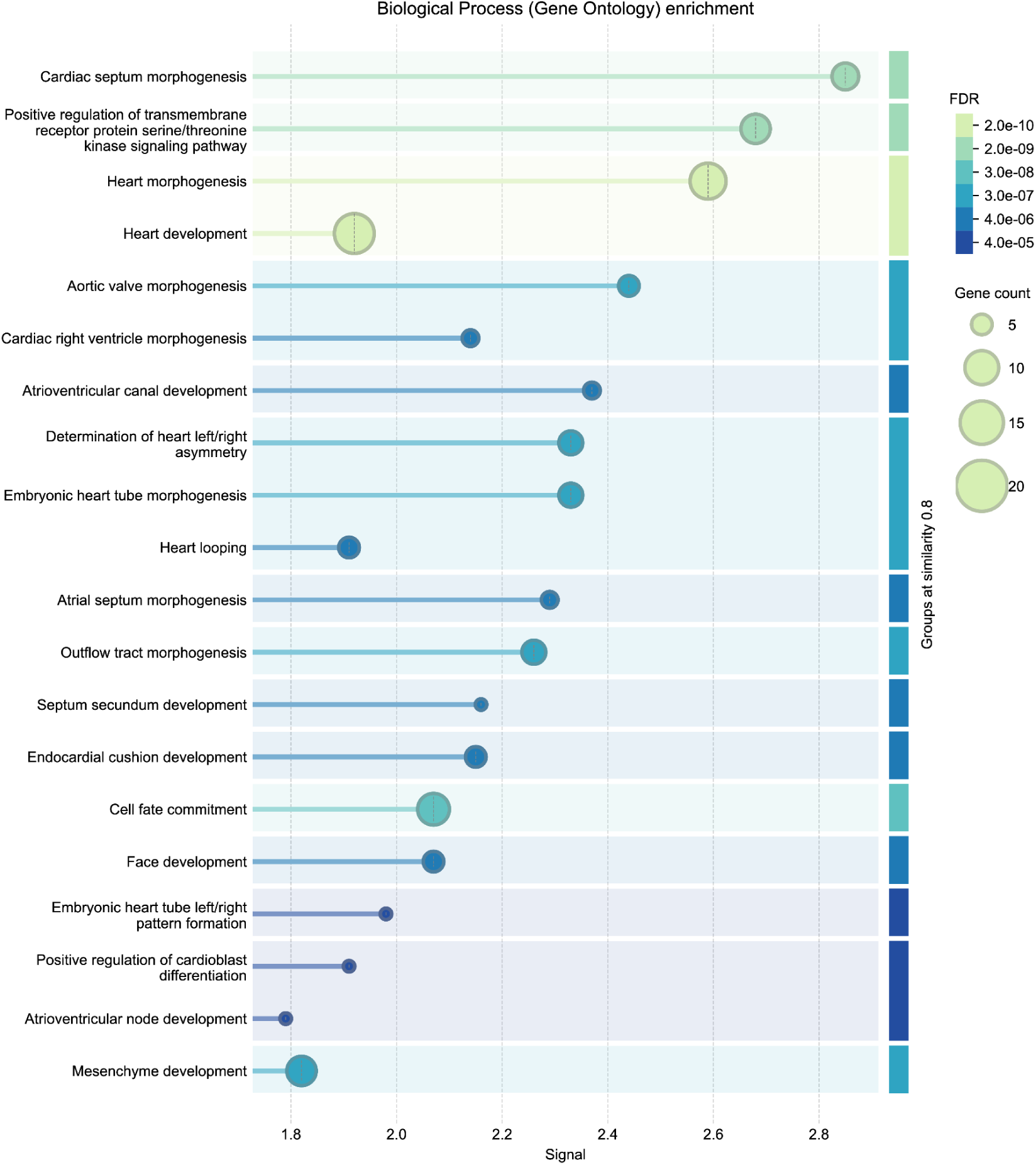
STRING enrichment analysis from Gene Ontology Biological Process for 27 genes identified from DDKG-based overlap between CHD cohort variants and IMPC phenotype-matched orthologs (Method 7). Results show significant overrepresentation in the 27 genes for developmental and signaling processes including different categories of cardiac development. FDR-adjusted p-values and gene set sizes are visualized using bar length and circle size, respectively. These results independently validate the biological relevance of DDKG-derived gene prioritizations in a cross-species phenotype-matching context.

**Figure 6.**
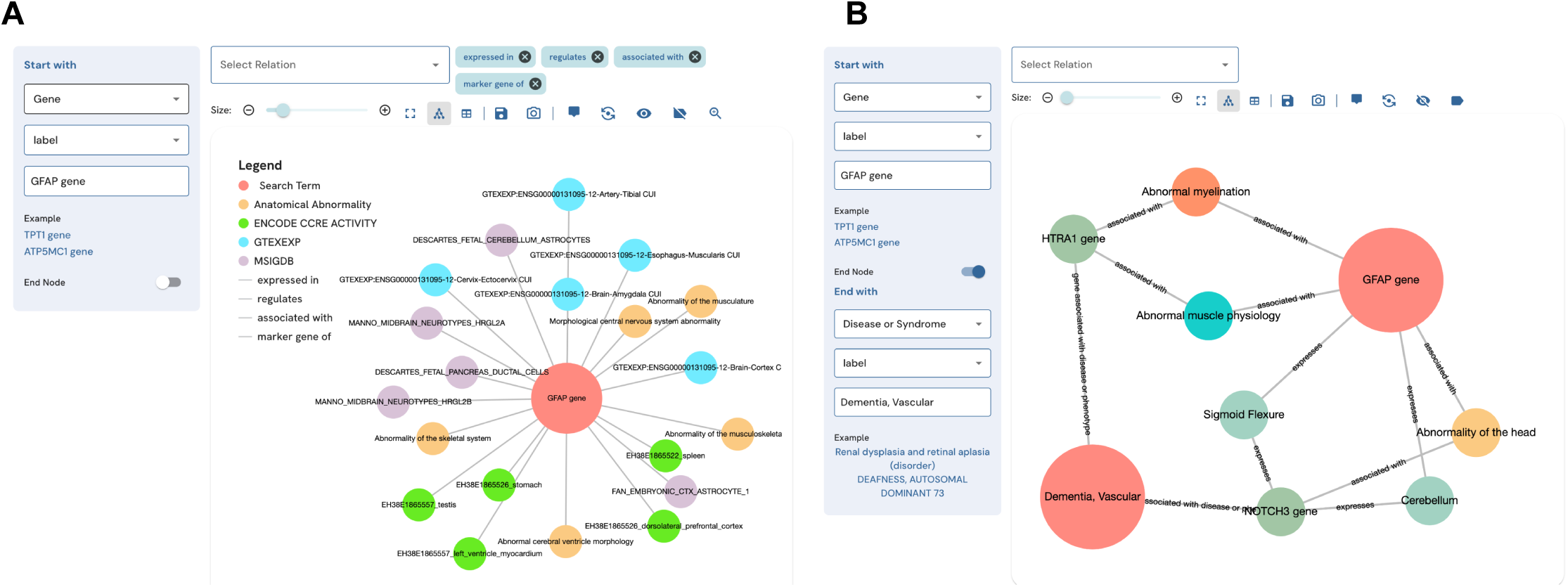
Visualization of gene-phenotype-disease associations using the DDKG-UI. The DDKG-UI, an interactive web-based platform derived from the Data Distillery Knowledge Graph (DDKG), enables users to explore complex biological relationships through customizable queries. (**A**) A query for the GFAP gene returns its expression across multiple tissues, anatomical abnormalities, and functional annotations. The graph illustrates associations with GTEx expression data, ENCODE regulatory elements, MSIGDB gene sets, and disease-related ontologies. (**B**) A disease-focused query links GFAP to vascular dementia, showing intermediary relationships with other genes (e.g., HTRA1 and NOTCH3) and relevant phenotypes, including abnormal myelination and muscle physiology alterations. The DDKG-UI allows researchers to dynamically search, filter, and visualize multi-omic datasets, facilitating hypothesis generation and biomedical discovery.

The analysis shows that brain tissues displayed a distinctive glycosylation profile, including FUT9, a known neural-specific α1,3-fucosyltransferase essential for Lewis X glycan formation involved with neurite outgrowth^35^. Another gene, MGAT3, is broadly expressed across neural tissues and is known to contribute to brain-specific bisecting GlcNAc glycan synthesis, as suggested by elevated HNK-1 glycan levels in Mgat3-deficient mouse brain^36^. Several other examples of tissue-specific glycoenzyme expression profiles from the integration of GlyGen and GTEx data provide a potentially rich field of research directions.

A single DDKG query was able to source the data and avoided manual data extraction. The same approach can be applied to any gene or other chromosomal feature to support reproducible, expression-driven exploration of functional gene sets across tissues and conditions.

### 2.4 Method 4: Identifying Genes and Associated Drug Targets

This method illustrates how the DDKG performs native multi-hop traversal across multiple biomedical data types. By connecting disease phenotypes to compounds via gene regulation and bioactivity data, it demonstrates how semantically structured graph queries can integrate molecular, expression, and pharmacological knowledge in a single step, without time-intensive manual reconciliation across resources.

Traditional in-silico drug discovery analyses often require manually linking disease-associated genes to perturbation data, target annotations, and chemical compounds across disparate sources. The DDKG encodes these relationships in a harmonized property graph, enabling direct traversal from phenotype terms to candidate drug targets and associated compounds. This reduces the complexity of cross-database queries and allows for interpretable, scalable exploration of disease mechanisms and therapeutic opportunities. The DDKG and its supporting open-source software packages are constructed to allow for rapid incorporation of new data, allowing users to seamlessly incorporate proprietary data into the graph for custom drug discovery pipelines.

This use case focuses on showing how to link diseases to genes to therapeutic compounds. Here we chose to examine the lipoxygenase gene family and its role in asthma-associated inflammation to identify potential therapeutic compounds. Inflammation causes airway swelling and obstruction ^37,38^, and human ALOX5 encodes a leukotriene-producing enzyme expressed in immune and epithelial cells. Leukotrienes are potent drivers of allergic and inflammatory disease^39,40^, including asthma^41^.

We used **Query 5-7** (Online Methods) to connect ALOX genes to asthma phenotypes using integrated data from IDG–DrugCentral, LINCS, and GTEx. ALOX5 was found to be negatively regulated by 39 compounds and positively by four ^42,43^ (Online Methods, **Figure 9** and **Figure 10**). Additional queries retrieved associations for ALOX15, ALOX15B, and ALOX12 (**Figure 11**), including the lipoxygenase inhibitor MLS000536924 (PUBCHEM:1778842) linked to ALOX15B ^43^.

**Figure 7:**
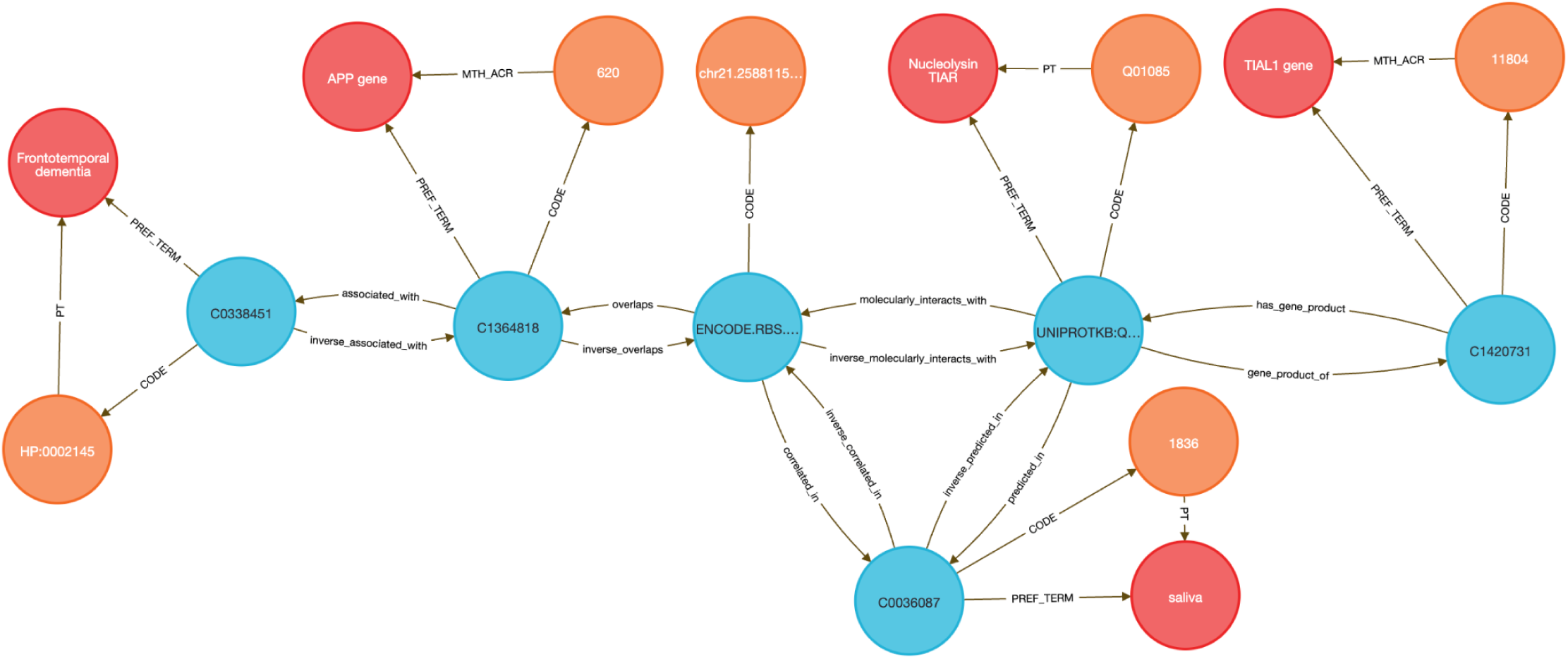
Example output from Query 2, illustrating a targeted liquid biopsy workflow for Frontotemporal Dementia (HP:0002145) by leveraging exRNA detection in saliva. The visualization highlights key molecular interactions between disease-associated genes, RNA-binding proteins (RBPs), and extracellular RNA (exRNA) expression patterns across relevant biofluids. The DDKG query framework identifies genes linked to Frontotemporal Dementia, the biofluids where their exRNA is detected, and the RBPs predicted to interact with those exRNAs, enabling insights into disease biomarker discovery. This structured graph-based approach facilitates hypothesis generation, in this case for non-invasive biomarker detection in neurodegenerative disorders.

**Figure 8.**
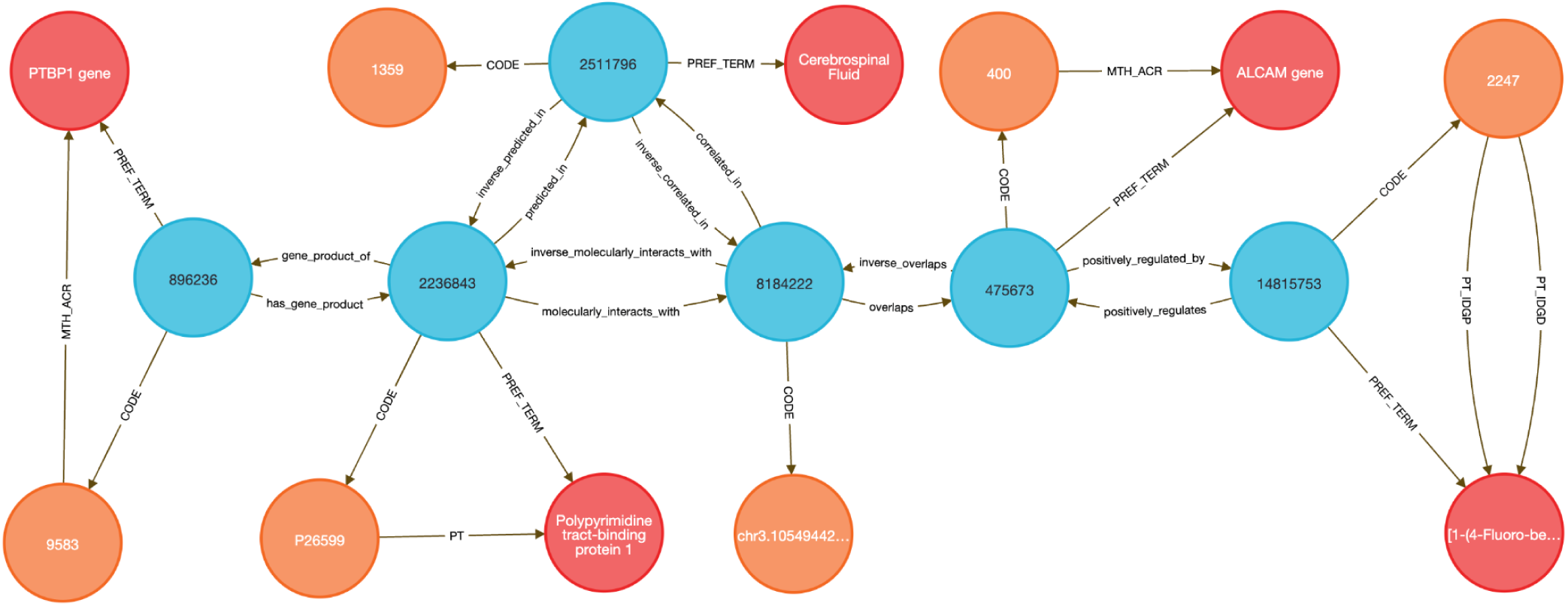
Liquid Biopsy Analysis. Approach for monitoring drug response to Astemizole, focusing on ALCAM expression and PTBP1 detection in cerebrospinal fluid. This subgraph is a result from the Query 3 path connecting the antihistamine Astemizole (PUBCHEM:2247) to its transcriptomic targets, specifically highlighting the ALCAM gene. ALCAM is positively regulated in LINCS data following Astemizole perturbation. The ALCAM locus overlaps an exRNA region bound by the RNA-binding protein PTBP1. PTBP1 is computationally predicted to be present in cerebrospinal fluid (CSF) and interacts with the overlapping exRNA locus, making it accessible via CSF pulldown. This example demonstrates how the DDKG enables integration of pharmacogenomic, regulatory, and tissue-localization data to support hypothesis generation for drug monitoring via targeted liquid biopsy.

**Figure 9:**
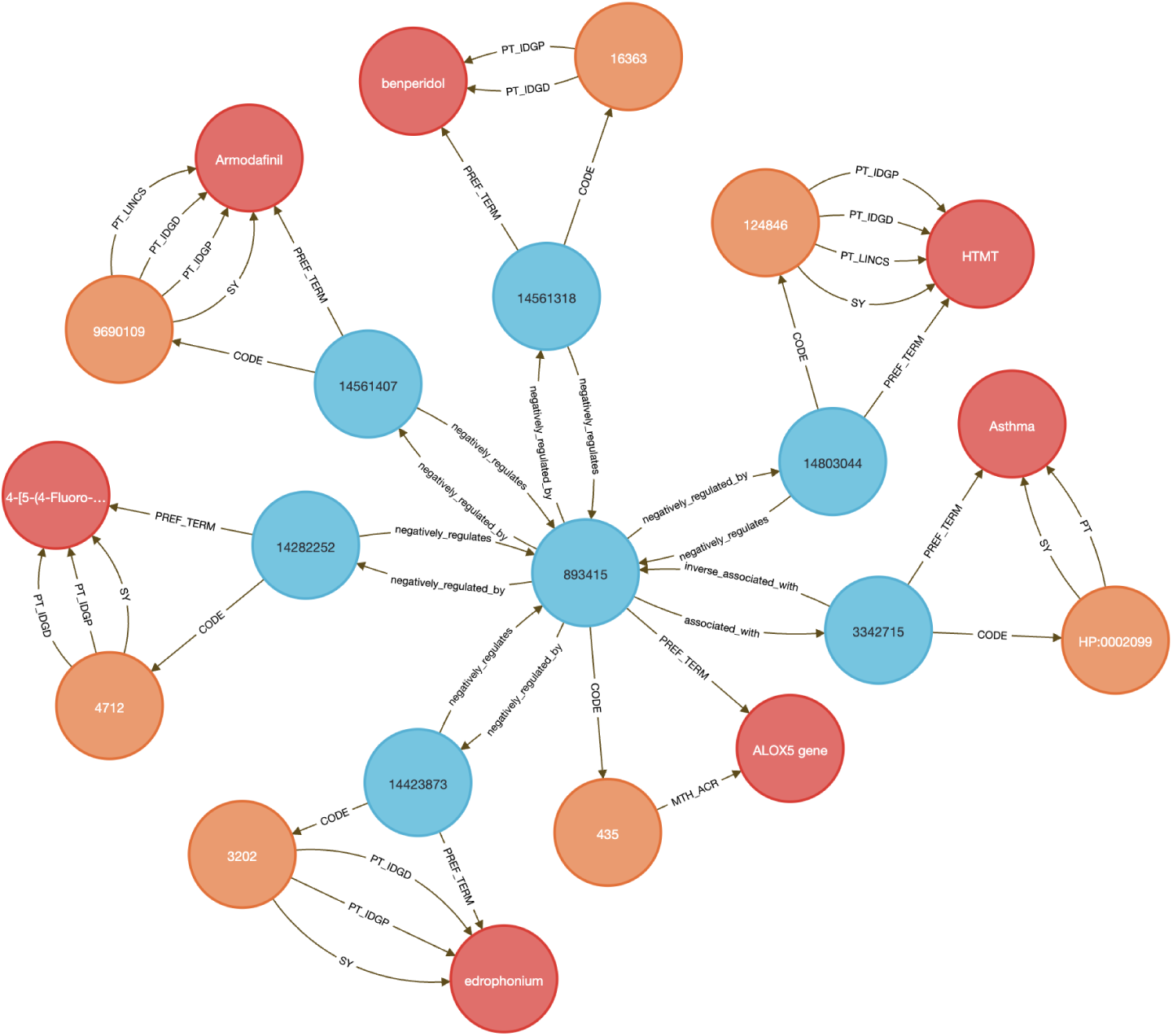
Combining data from IDG and LINCS, shown are the results for Query 5 seeking genes and Pubchem compounds associated with the disease “Asthma.”

**Figure 10:**
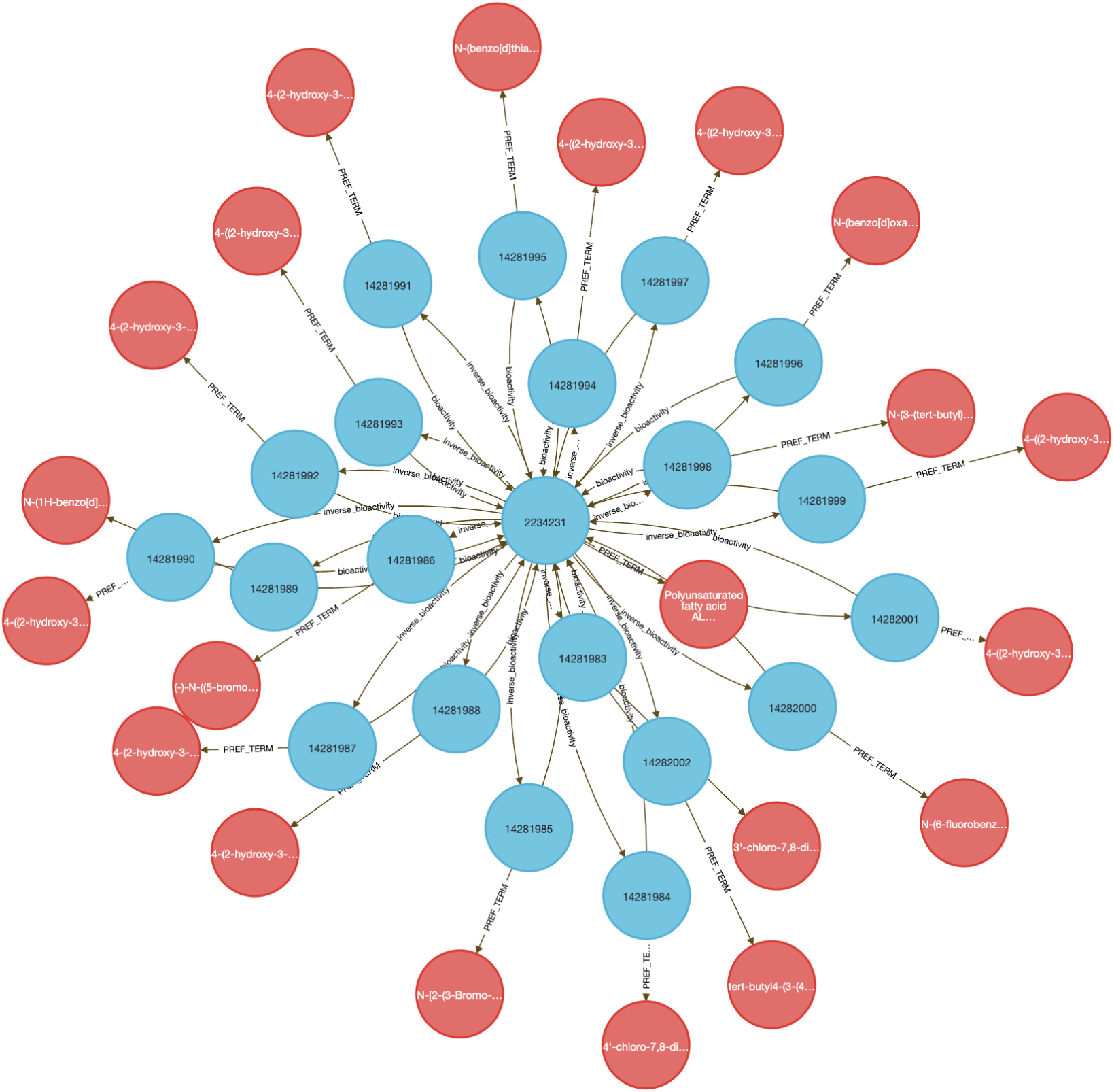
Cypher query results for genes containing the string “ALOX” (Query 6). This query returns human lipoxygenase genes along with Pubchem compounds associated via the bioactivity relationship.

**Figure 11:**
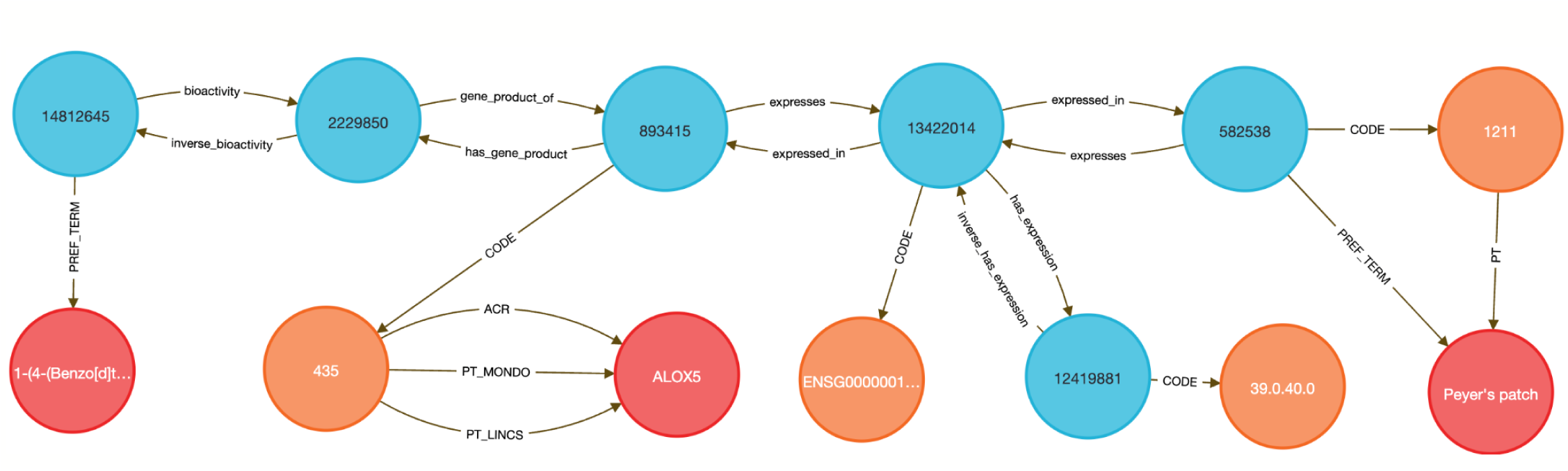
Graph-based representation of ALOX5 compound activity and expression profile. This example subgraph is one result from Query 7. It shows the connection from a PubChem compound with known bioactivity (from IDG) to its UniProt gene product, the HGNC-coded gene ALOX5. GTEx expression data is linked through expression bins and mapped to UBERON tissues (here Peyer’s patch). The result demonstrates the DDKG’s integration of chemical, molecular, and anatomical data, enabling rapid exploration of tissue-specific gene–compound interactions.

Tissue-specific expression analysis confirmed ALOX5 activity in breast epithelium, Peyer’s patches, and the anterior lingual gland, consistent with prior reports ^44–47^, validating the biological plausibility of the DDKG-derived results and hypothesis of inhibitors like MLS000536924 (**Figure 11**).

While this case centers on asthma and lipoxygenases, the same approach can be generalized to drug repurposing, compound prioritization, and tissue-aware target validation across disease contexts. All results were retrieved in a single graph-native query, demonstrating how integrating DDKG datasets across

IDG-derived knowledge can drive rapid, reproducible therapeutic discovery.

### 2.5 Method 5: Integrating Metabolite Related Studies

Integrative metabolomics remains a major challenge due to the need to harmonize drug bioactivity, gene regulation, metabolite profiles, tissue expression, and disease phenotypes—data types often siloed in distinct repositories with incompatible identifiers. Analysts typically rely on custom pipelines to align pathway, expression, and clinical data. The DDKG simplifies this process by enabling seamless traversal across semantically linked pharmacologic, genomic, and metabolomic entities. While this use case focuses on compound-mediated metabolic relationships, the framework supports any multi-modal systems biology analysis.

Using data from Metabolomics Workbench (MW), IDG–DrugCentral, and GTEx, we used **Query 8** to trace paths connecting compounds, protein targets, encoding genes, downstream metabolites, associated disease phenotypes, and relevant tissues (**Figure 4**). This enabled identification of molecular intermediates linking drug action to disease via metabolic signatures.

The query returned over 450,000 associations, filtered to retain biologically plausible connections in well-annotated tissues and disease contexts. Node-level co-occurrence analysis highlighted genes such as CYP3A4 and TP53, and metabolites involved in inflammatory and detoxification pathways ^15,24^ .

Performing this analysis using typical bioinformatics approaches would require manual joins across at least four distinct data sources. In contrast, the DDKG query was executed in seconds and returned reproducible, exportable results, accessible via both Neo4j and the public DDKG-UI.

This use case demonstrates how the DDKG facilitates systems-level exploration of pharmacogenomic and metabolic pathways, accelerating hypothesis generation across molecular, cellular, and phenotypic contexts.

### 2.6 Method 6: Cross-Species Integration of Gene Expression and Regulatory Variation

Linking transcriptional data from animal models to human regulatory mechanisms is a longstanding challenge in systems genomics. Traditional approaches require orthology mapping, data harmonization, and annotation pipelines that involve extensive scripting and manual curation. The DDKG simplifies this process through semantically harmonized, query-driven integration across species and data types.

We examined whether genes responsive to exercise in a rat model are also under genetic regulation in the human heart. Differentially expressed genes from the MoTrPAC rat heart dataset were mapped to human orthologs using HGNC’s Comparison of Orthology Predictions (HCOP, ^48^. The top 500 human genes were selected based on GTEx heart eQTL counts, reasoning that higher regulatory burden may correspond to exercise-induced variability. A single DDKG query returned all gene–variant–tissue associations linking the rat response to human regulatory variation (Online Methods **Query 9**, **Figure 12**).

**Figure 12:**
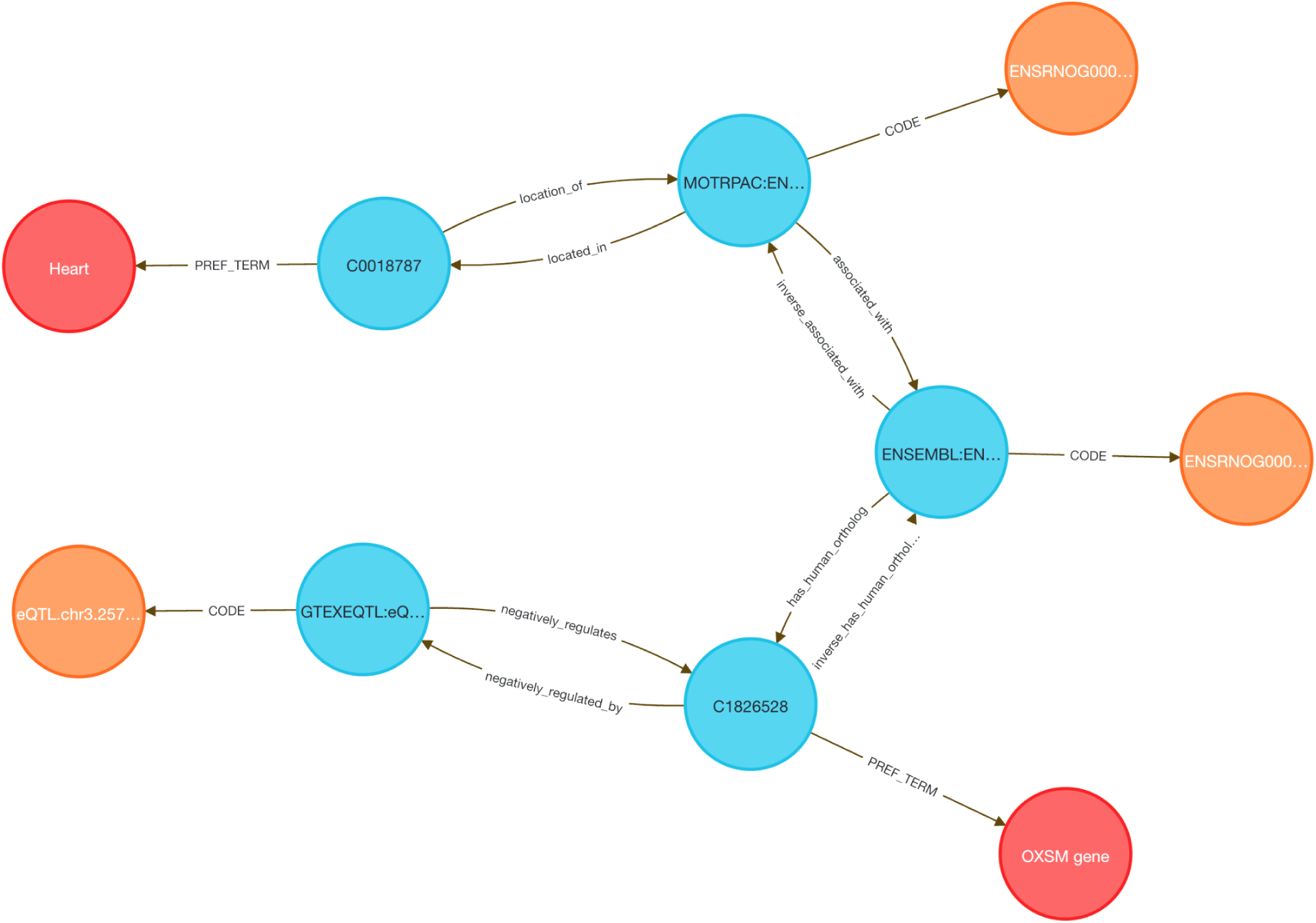
Query 9 aims to identify data points that are linked to evidence related to genes in tissues across different omes in the MoTrPAC young adult rats endurance training exercise data that match GTEx eQTL regulation in heart. The resulting gene set are exercise-linked genes with tissue expression matches to humans that are linked to disease.

To contextualize these results, we applied MSigDB pathway enrichment (Hallmark, C2.CP, HPO), identifying sets related to oxidative phosphorylation, mitochondrial translation, and respiratory electron transport—hallmarks of cardiovascular adaptation to exercise.

Several genes have plausible mechanistic roles. PIEZO1, a mechanosensitive calcium channel, regulates stretch-induced ROS signaling and is linked to cardiomyopathy and heart failure in mouse and human models ^49,50^. GNA12 regulates Rho-GTPase signaling and contributes to vascular remodeling. CD151, a tetraspanin that complexes with integrins such as α6β1, promotes post-ischemic angiogenesis via PI3K/Akt/eNOS and FAK signaling ^51^, and is inducible by mechanical stress ^52,53^, consistent with exercise-induced vascular remodeling and its high eQTL count in GTEx. Other genes such as BSG (CD147) and HLA-A/C are involved in immune and matrix remodeling, while FLCN has been shown to regulate cardiac hypertrophy via mTORC1 signaling in murine models^54^. Additional genes in the set could be hypothesized to contribute to metabolic and mechanical stress adaptation pathways.

This use case illustrates how the DDKG enables rapid, reproducible integration of cross-species omics and regulatory data using a single graph-native query.

### 2.7 Method 7: Semantic Graph Querying for Cross-Species Variant–to-Phenotype Prioritization

Linking rare disease variants to model organism phenotypes is a widely used strategy for inferring gene function, especially when clinical studies lack functional follow-up. Traditional workflows require orthology mapping, phenotype filtering, and harmonization of patient variants, gene annotations, and disease ontologies—often through complex, multi-step scripting. The DDKG streamlines this process by enabling a single semantically structured query that integrates these data sources to support translational insights into gene function and disease mechanisms.

In this use case, we analyzed germline variants from a pediatric congenital heart defect (CHD) cohort (dbGAP phs001138.v4.p2) using data from the Kids First Data Resource Center (KFDRC). Variants were binned by gene (see Online Methods) and mapped to mouse orthologs via HGNC’s Comparison of Orthology Predictions (HCOP) ^48^, then to phenotypes from the International Mouse Phenotyping Consortium (IMPC) ^55^, including data from the NIH Common Fund Knockout Mouse Phenotyping Project, KOMP2. A single DDKG query identified orthologous mouse genes with phenotypes matching those observed in the CHD cohort. We focused on atrial septal defect (ASD) as a representative phenotype.

The integrated query (Online Methods **Query 10**) completed in seconds and returned 27 genes including known cardiac developmental regulators, including GATA4, GATA5, GATA6, NKX2-5, TBX20, CITED2, CREBBP, and EP300, all with established roles in septation, chamber specification, and congenital heart defects ^56–62^ . TBX20, for example, is essential for chamber formation; murine knockouts show severe cardiac malformations, while gain-of-function variants in humans are linked to ASD and patent foramen ovale ^59,63,64^. Additional top-ranked genes included NSD2, an epigenetic regulator implicated in pulmonary arterial hypertension^65^ and RAS/MAPK pathway genes BRAF, RAF1, and PTPN11, which drive Noonan syndrome and other RASopathies associated with CHD. GDF1 and NODAL, regulators of left–right patterning and outflow tract development, were also recovered in the query ^66,67^ . The list of 27 genes can be found in **Supplementary File 2.**

To independently validate the biological coherence of these results, we submitted the 27 ASD-linked genes to STRING (v12.0) ^68^ for enrichment analysis using the Gene Ontology Biological Process category ^69,70^. The results (**Figure 5**) yielded results related to several categories of cardiac development, with FDR values below 1e-5. The external analysis confirms that the DDKG query prioritized biologically plausible candidates, and demonstrates the utility of semantic graph-based phenotype matching for rapid and reproducible cross-species gene prioritization.

This use case illustrates how the DDKG supports hypothesis-driven exploration of human variant function across species by embedding cross-domain annotations in a single graph-native framework. The approach is extensible to other cohorts and phenotypes and can be generalized for use in translational gene discovery workflows.

### 2.8 Method 8: Semantic Graph Querying for Cross-Disease Gene Discovery

Identifying shared molecular mechanisms across distinct disease categories remains a major challenge in translational genomics research. This use case illustrates an approach to the problem by asking whether congenital heart defects and childhood leukemias might share common gene drivers that are detectable through graph-based similarity measures. Decades of previous work has established epidemiological evidence for links between birth defects and pediatric cancer incidence, one recent study surveyed millions of pediatric records ^71^. We selected two phenotypes representing congenital heart disease (Tetralogy of Fallot, HP:0001636; Atrial Septal Defect, HP:0001631) and two representing pediatric leukemia (Acute Lymphoblastic Leukemia, HP:0006721; Acute Myeloid Leukemia, HP:0004808), and used the DDKG to search for human genes positioned closest to all four identifiers within the graph topology.

**Query 11** (Online Methods) applies the Neo4j Graph Data Science (GDS) library’s Common Neighbors (CN) algorithm to compute similarity scores between each phenotype and all protein-coding genes in the graph.

Disease-Phenotype–Gene connections in the DDKG are derived from integrated associations from sources like HPO, ClinVar, OMIM, and Orphanet. For each phenotype, genes were ranked by their CN score (higher scores indicating greater overlap in graph neighborhood). We then summed the ranks for each gene across all phenotypes and re-ranked genes by their aggregate rank, yielding a prioritized list of candidates most likely to be implicated in both congenital and hematologic disease contexts (see Online Methods).

The resulting thirty top-ranked genes (**Table 1**) include canonical regulators such as *BRAF*, *NF1*, *KRAS*, *PTPN11*, and *TP53* which have known roles in both developmental syndromes and oncogenic signaling. To evaluate the biological relevance of the top-ranked genes from the CN analysis, we performed independent enrichment analysis at the STRING website’s “search” function (at https://string-db.org)^68^. We submitted the top 200 ranked genes from our analysis as described in Online Methods to STRING’s functional annotation tool. Results showed strong signals for Human Phenotype Ontology terms related to cancer and developmental defects, consistent with the input phenotypes. Notably, the top enriched HPO terms included Leukemia and Atrial septal defect, with FDR values below 1e-52 (see **Figure 15** in Online Methods). These results provide orthogonal validation that the DDKG-derived rankings calculated from **Query 11** results reflect biologically relevant gene-phenotype relationships and can provide hypotheses for shared mechanisms across disease categories. The full table of ranked results including the first thirty in **Table 1**, are given in **Supplementary File 3**.

**Figure 13:**
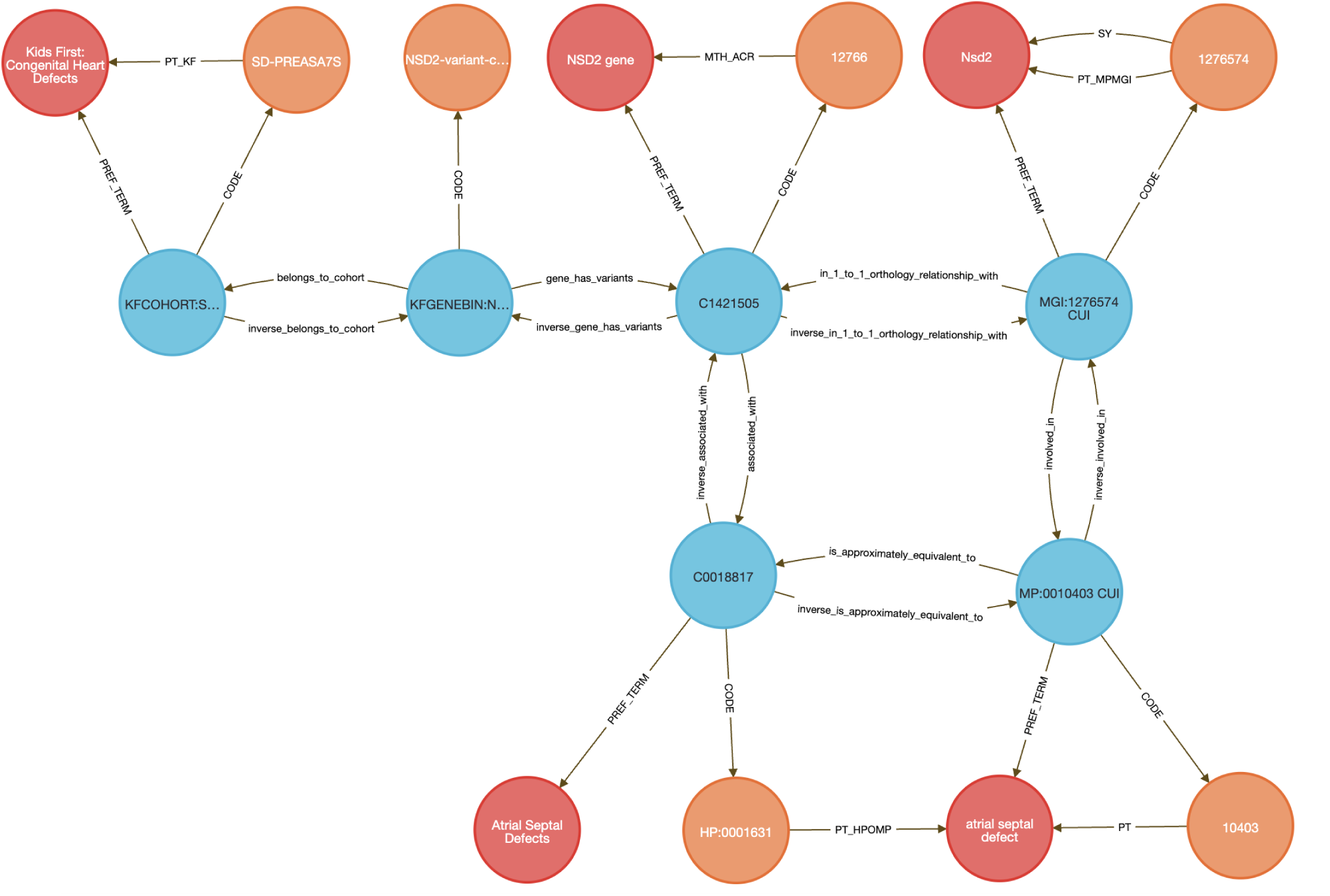
Cross-species integration of mouse phenotypes with human genomic data using the DDKG. Mouse phenotypes associated with atrial septal defects (left) are linked to IMPC-derived gene knockouts, then mapped to human orthologs via HCOP. These human genes are associated with pathogenic variants in the Kids First congenital heart defect cohort (right). This multi-ontology traversal demonstrates how model organism data can inform human disease gene discovery through knowledge graph integration.

**Figure 14.**
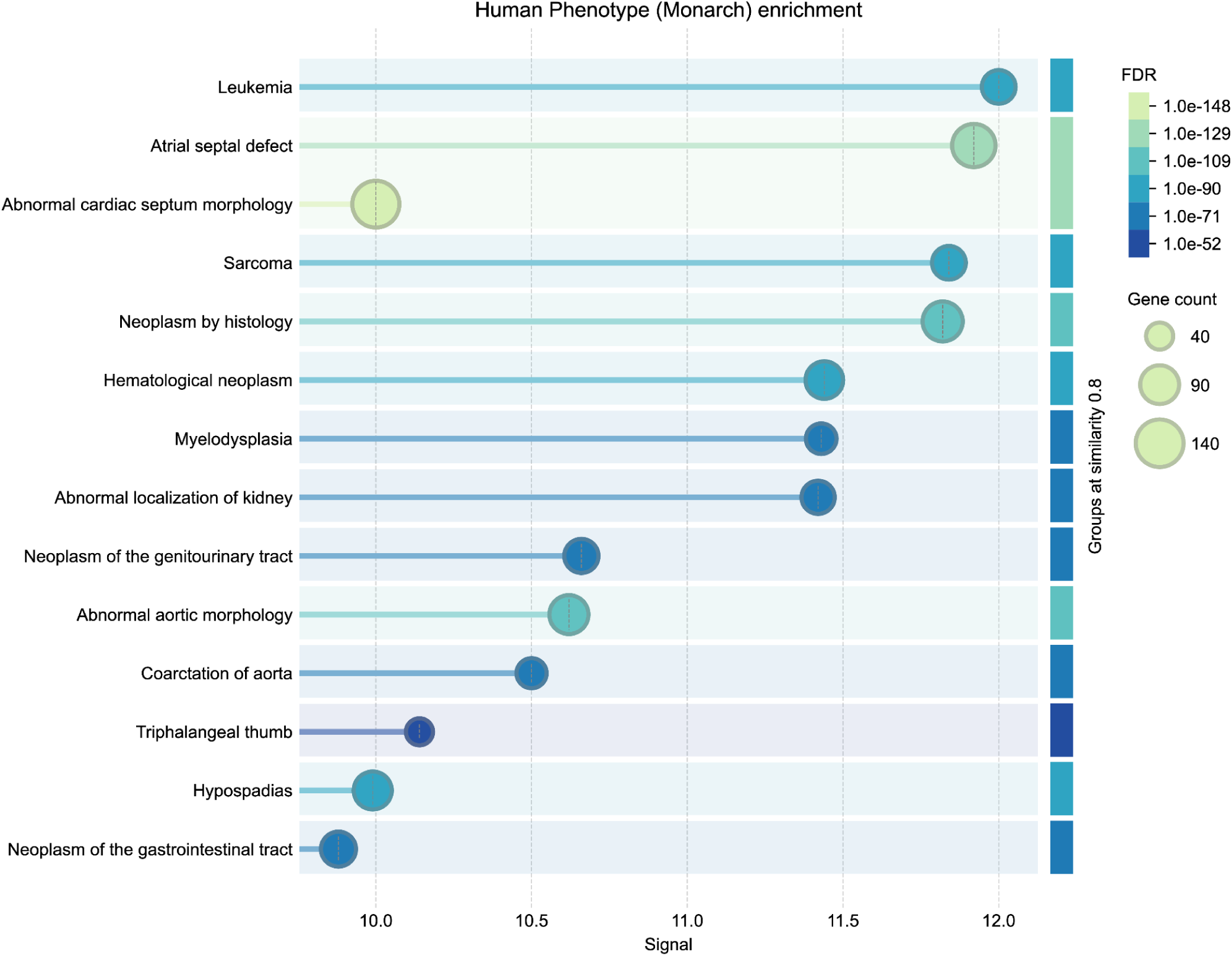
STRING gene enrichment results on the Human Phenotype Ontology for the top 200 genes ranked by DDKG graph-based proximity to congenital heart defects and pediatric leukemias (Method 8). Genes prioritized using the Common Neighbors algorithm (**Query 11**) were analyzed using the STRING v12.0 functional annotation tool. The plot shows statistically significant signals for Leukemia and Atrial Septal Defects within the Human Phenotype Ontology (HPO) category.

**Figure 15.**
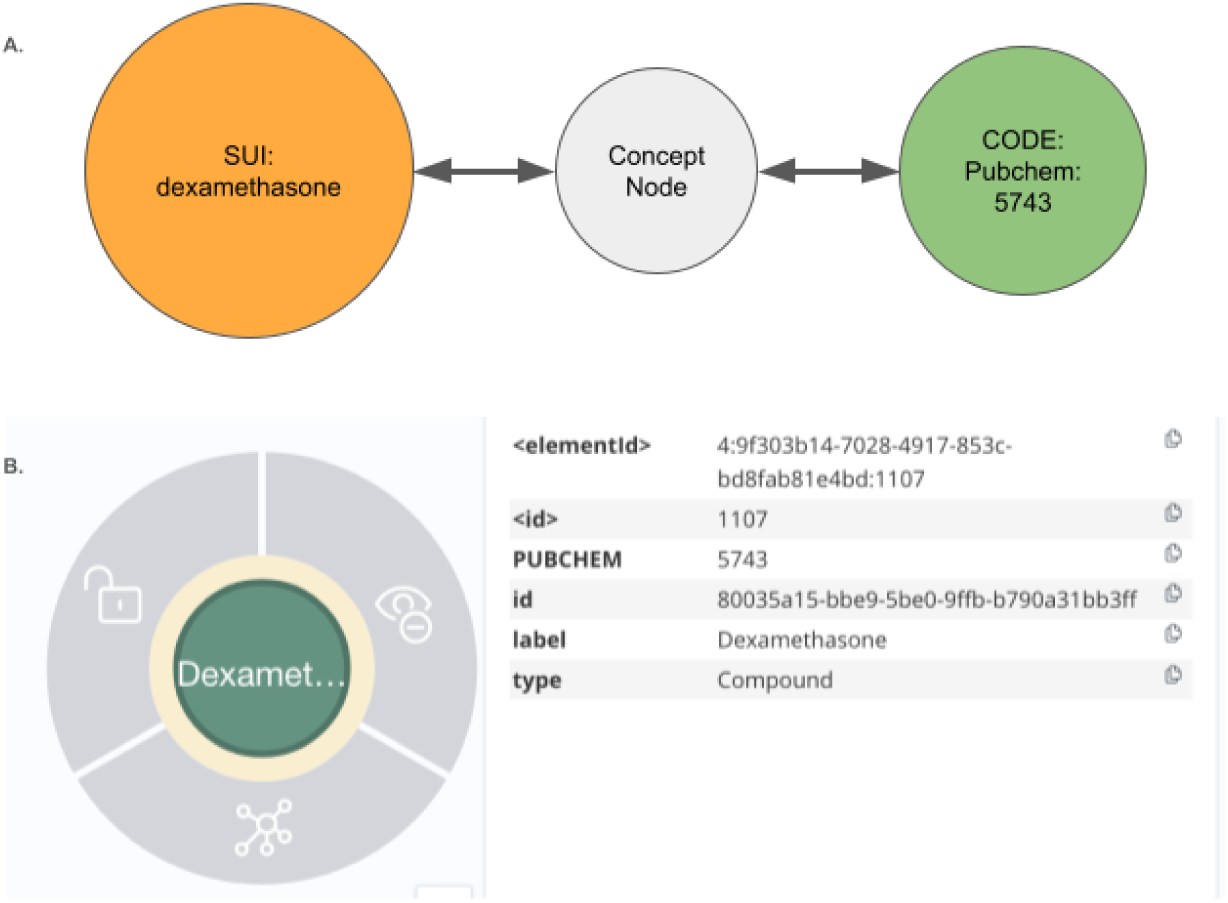
Converting UBKG assertions into DDKG-UI assertions. The DDKG has limited properties on the Concept nodes. In this figure we show an example of a transformed node by extracting Code and Term information into properties on the concept nodes in the DDKG-UI subgraph.

**Table 1:** First 30 top-ranked genes by proximity to phenotypes spanning congenital heart disease and pediatric leukemia. Numbers in columns indicate the rankings (1->N) of common neighbor scores for that phenotype. Genes were prioritized using the Common Neighbors (CN) algorithm from the Neo4j Graph Data Science library, based on their topological similarity to four selected Human Phenotype Ontology (HPO) Terms: ToF: Tetralogy of Fallot (HP:0001636), ASD: Atrial Septal Defect (HP:0001631), ALL: Acute Lymphoblastic Leukemia (HP:0006721), and AML: Acute Myeloid Leukemia (HP:0004808). For each phenotype, genes were ranked by CN score, and ranks were summed across all phenotypes to produce a composite rank, ordered here. Genes with the lowest total ranks are therefore those most proximal to all four phenotypes in the DDKG graph topology. The thirty top-ranked genes are enriched for developmental and oncogenic pathways (see **Figure 13** in Online Methods).

| HPO ID -> | HP:0001636 | HP:0001631 | HP:0006721 | HP:0004808 |
| --- | --- | --- | --- | --- |
| Gene_Symbol | ToF | ASD | ALL | AML |
| BRAF | 38 | 30 | 5 | 7 |
| NOTCH1 | 8 | 25 | 72 | 58 |
| NF1 | 94 | 35 | 46 | 35 |
| KRAS | 183 | 6 | 15 | 9 |
| HRAS | 193 | 15 | 24 | 18 |
| GATA1 | 110 | 39 | 47 | 55 |
| RPL5 | 57 | 31 | 45 | 121 |
| PTPN11 | 192 | 50 | 16 | 12 |
| MAP2K1 | 215 | 16 | 44 | 54 |
| FGFR2 | 58 | 32 | 144 | 140 |
| KMT2A | 213 | 139 | 7 | 16 |
| FGFR1 | 95 | 36 | 145 | 102 |
| ASXL1 | 216 | 141 | 8 | 28 |
| SETBP1 | 214 | 140 | 26 | 20 |
| CREBBP | 247 | 55 | 74 | 59 |
| NSD1 | 217 | 142 | 48 | 36 |
| BCR | 248 | 147 | 1 | 47 |
| MAPK1 | 196 | 51 | 73 | 126 |
| ABL1 | 275 | 156 | 17 | 10 |
| EP300 | 249 | 56 | 75 | 88 |
| NOTCH2 | 26 | 85 | 148 | 225 |
| WT1 | 41 | 277 | 83 | 91 |
| PALB2 | 96 | 289 | 85 | 92 |
| SETD2 | 250 | 148 | 78 | 89 |
| BRCA2 | 194 | 320 | 27 | 32 |
| TP53 | 191 | 404 | 2 | 2 |
| FANCE | 49 | 279 | 84 | 205 |
| GPC3 | 179 | 314 | 87 | 60 |
| FLCN | 338 | 61 | 146 | 103 |
| TGFBR2 | 251 | 149 | 150 | 104 |

**Table 2.**
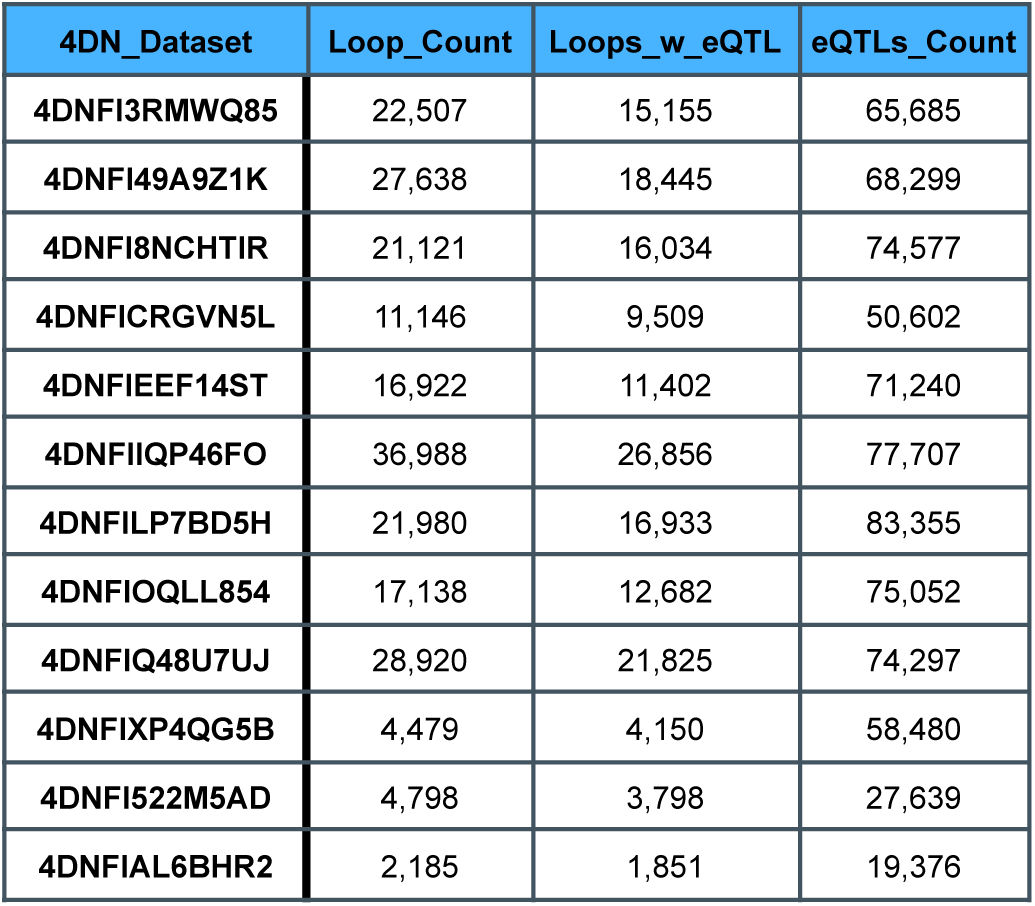
Summary statistics for 4DN chromatin loop datasets analyzed in Method 1. Each row represents a dataset from the 4D Nucleome (4DN) project loaded into the DDKG. Columns report the total number of loops (Loop_Count), the number of loops overlapping at least one GTEx eQTL (Loops_w_eQTL), and the total number of GTEx eQTLs found within those loops (eQTLs_Count). These results were obtained using the HSCLO structure at 1 kb resolution, demonstrating the prevalence of regulatory variant overlap with 3D genomic features across multiple cell types and experimental conditions. The aggregated data form the basis for topological analyses of regulatory architecture using the DDKG.

**Table 3:** UniprotKB IDs for each Lipoxygenase isoform.

| UNIPROTKB | Lipoxygenase isoform |
| --- | --- |
| O15296 | *ALOX15B* |
| P16050 | *ALOX15* |
| P18054 | *ALOX12* |

### 2.9 The DDKG-UI: Web Interface to the DDKG

The DDKG was designed not only to support use by computational experts, but also to be accessible to researchers with no prior experience in graph manipulation. The DDKG-UI was developed to provide a web-based interface for intuitive search and exploration, with guided query templates and prebuilt use case “apps” for common biomedical questions, such as tissue-specific gene expression, regulatory variant analysis, or biomarker discovery in liquid biopsies. These features help ensure that users at all levels can engage with the graph in biologically meaningful ways, reducing the risk of nonsensical queries and supporting reproducible, interpretable analyses. The DDKG-UI can be accessed at https://dd-kg-ui.cfde.cloud/

We provide an example of querying the DDKG-UI for Common Fund datasets to gain knowledge about the gene GFAP (**Figure 6**). GFAP is a gene that is primarily expressed in the central nervous system. Plasma levels of GFAP have been linked to several nervous system disorders ^72^. A recent breakthrough study has identified GFAP as a potential biomarker for dementia, with changes in its expression observed at least ten years prior to the onset of the disease ^73^. The DDKG-UI single search term returns immediate neighbors of the gene GFAP (**Figure 6a**). We observe that GFAP is linked to several tissues and cell types of the central nervous system as well as its association with abnormalities of the central nervous system. Next, we look at the connections between GFAP and Vascular Dementia (VD) (**Figure 6b**). We see that GFAP is connected to VD via another gene that is also associated with abnormal myelination. It has been documented that myelin dysfunction is a driving factor for amyloid β (Aβ) deposition in Alzheimer’s disease ^74^. Aβ is known to be deposited in cerebral blood vessels and has been linked with vascular cognitive impairment ^75^.

## 3. Discussion

Knowledge graphs provide a scalable framework for integrating and analyzing structured biomedical data. They enable exploratory analysis across molecular, clinical, and phenotypic domains within a unified semantic structure. Effective use of such graphs depends on the quality of the data they represent. Expert annotation ensures that relationships in the graph reflect accurate and domain-relevant knowledge. The DDKG integrates curated datasets from NIH Common Fund Data Coordination Centers alongside public resources such as ClinVar and STRING, embedding high-confidence annotations where possible and flagging community-contributed data where standards may vary. This approach supports reliable cross-domain exploration and reuse.

The DDKG was developed within the NIH Common Fund Data Ecosystem (CFDE) to advance interoperability and reuse across projects including 4DN, GTEx, Metabolomics Workbench, GlyGen, HuBMAP, LINCS, and Kids First. These programs have produced multi-modal datasets from diverse cohorts and experimental platforms that can be further integrated to address many biomedical research questions. The DDKG helps with this integration by embedding these resources into a highly connected property graph built on the UMLS Concept–Code–Term model. It incorporates over 180 ontologies and vocabularies, aligning Common Fund data with additional datasets including ENCODE, GENCODE, ENSEMBL, and STRING. Relationships among genes, variants, tissues, phenotypes, compounds, and diseases are encoded as typed edges across semantically normalized nodes. The DDKG requires an UMLS license to download, as the DDKG includes the UMLS within the build.

Rather than layering annotations after graph construction, the DDKG integrates data at the point of ingestion using scripts that apply ontological mapping during the loading process. This method ensures that biological meaning is preserved and made consistent across sources. It also avoids redundant workflows by enabling direct multi-hop queries across molecular and clinical data types without manual joins.

The use cases highlight the range of bioinformatics and clinical informatics workflows supported by the DDKG. These include matrix-based expression analysis, regulatory annotation, phenotype linkage, target discovery, and cross-species inference. The graph structure enables hybrid queries that span clinical vocabularies and molecular features. **Method 1** integrated chromatin conformation with regulatory variants to recover enhancer–promoter spatial patterns. **Method 2** identified candidate biomarkers using exRNA and RNA-binding protein annotations. **Method 3** explored tissue-specific glycosylation signatures in GTEx. **Method 4** linked disease genes to drug perturbation profiles, recovering known targets such as ALOX5. **Method 5** reconstructed metabolite-associated pathways across genes, compounds, and disease phenotypes. **Method 6** matched exercise-induced gene expression in rats to human heart eQTLs. Each was executed with queries demonstrating that common analytical tasks can be reframed as graph traversals without significant manual work.

**Method 7** demonstrates how semantically structured graph queries can streamline variant-to-phenotype inference across species. Integrating human variant data from a cohort with congenital heart disorders with orthology mappings and model organism phenotypes enables the DDKG to deliver biologically meaningful connections between clinical findings and experimental evidence. The rapid recovery of known congenital heart disease genes such as TBX20^59,64^, GATA4^76^, and NKX2-5^58,76,77^, as well as RAS pathway-linked genes like PTPN11^78^ and RAF1^79^, shows that the DDKG effectively recapitulates core heart developmental mechanisms from human to mouse. This approach generalizes to other rare disease cohorts, offering a reusable workflow for cross-species inference that bypasses traditional data wrangling.

**Method 8** demonstrates how a calculation of semantic proximity in the DDKG can reveal both established and unexpected biology. We found several canonical regulators among the top-200 prioritized genes related to both congenital heart defects and pediatric leukemias, including BRAF, KRAS, PTPN11, TP53, and GATA1. These genes reflect pathways related to processes such as RASopathies and hematological malignancies.

As an example of DDKG queries driving new biological hypotheses, **Method 8** also ranks DLK1, a non-canonical NOTCH ligand not previously strongly associated with congenital heart defects, within the top 200 genes. DLK1 modulates NOTCH signaling during development ^80,81^ and expression changes have been linked to myelodysplastic syndromes and AML ^82^. A suggested annotation link between DLK1, cardiac phenotypes and leukemia could suggest another mechanism for a potential cross-disease role via NOTCH pathway modulation. This finding spotlights DLK1 as a hypothesis-driven candidate for future exploration for studying commonalities between hematological malignancies and congenital heart defects which have been established by previous epidemiological studies^71^. The prominence of RAS-pathway genes also helps confirm the method’s biological relevance, while identification of other genes like DLK1 can highlight the DDKG’s power to generate novel, biologically coherent avenues for new discovery.

The DDKG can also be extended with user-supplied datasets as it has been designed to be modular and extensible. New data, such as drug screening results, animal model experiments, patient records, single-cell markers, and additional omics datasets can all be integrated without modifying the schema if at least one new data identifier is already in the DDKG. Users can extend the graph locally by importing their files with the UBKG’s provided ingestion scripts, and/or by building the DDKG with desired modules, supporting customization and reuse.

Current limitations reflect considerations in performance and gaps in the underlying datasets. Currently the size of the DDKG will be limited by any platform requirements. Incorporating datasets from literature extraction can further the DDKG’s power for studying under-sourced domains such as rare diseases, environmental exposures, and epigenomic regulation. The DDKG updates through the UBKG bi-annual releases and will include ontology maintenance as community standards evolve. The DDKG-UI enables interactive access, although more comprehensive and advanced workflows will require direct interfacing with the DDKG and familiarity with graph query languages. Usability will improve with expanded APIs and visual tools as the resource matures over time.

The DDKG provides a reusable semantic graph that supports biomedical discovery across data types and contexts. Its structure promotes scalable integration and analysis without the need for custom data harmonization or repeated annotation, lowering barriers for multi-modal insight.

## 4. Online Methods

The application of knowledge graphs in biomedical research is growing rapidly, including the construction of networks from literature mining and other data sources linking genes, diseases, and drugs across curated databases. Yet, the power of knowledge graphs also introduces potential pitfalls where flexible graph queries may return technically correct results that are biologically irrelevant or misleading if posed without appropriate context. For example, a shortest-path query could trivially connect Alzheimer’s disease metabolites to pediatric cancer variants simply due to shared ontology terms, not because of any real biological relationship. Without expert oversight, the flexibility that makes these graphs powerful can also allow nonsensical or spurious conclusions.

To be effective, biomedical knowledge graphs must therefore be both constructed and used under the supervision of experts who understand the biological meaning of the data and the limitations of the underlying sources. Building a richly annotated biomedical knowledge graph is both an annotation and data engineering task, requiring the curation and alignment of relevant connecting entities that have been created by the biological community, such as GENCODE ^83^, Ensembl ^84^, UniProt ^85^, Reactome ^86^, WikiPathways ^87^, STRINGdb ^68^. Equally important is the application of biological judgment in deciding when and how entities should be linked, ensuring that relationships represent true biological insight rather than incidental data proximity. Without this domain knowledge, even the best-engineered graph architectures risk misrepresenting biological systems and misguiding downstream analyses.

Importantly, the DDKG, like Petagraph^10,11^, operates as a general-purpose knowledge graph data warehouse, optimized for data harmonization and query performance but is not pre-configured for machine learning because of its generalist nature. Consistent with Petagraph’s design principles, the DDKG represents many node attributes, such as identifiers, preferred terms, and metadata as separate, linked nodes rather than collapsed node properties. This schema promotes interoperability and flexible ontology alignment but differs from the feature-vector organization typically required for machine learning tasks. Preparing the DDKG for graph machine learning, such as node classification or link prediction, therefore requires subgraph projection of the desired data categories with repacking of these distributed properties onto parent nodes. This process must be tailored to a specific machine learning use case and we provide example scripts in our repository to assist. This separation of concerns between data integration and analytic modeling allows researchers to optimize graph preparation for their chosen analysis strategy while benefiting from a shared, curated knowledge resource that spans many different domains of interest.

Here, we describe the design, construction, and validation of the DDKG. We present its ontology-driven integration strategies, query infrastructure, and machine learning capabilities, and demonstrate its use through several representative biomedical applications. Together, these elements position the DDKG as a reusable, extensible platform for multi-omics data integration and a model for constructing biologically grounded knowledge graphs that support hypothesis generation, exploration, and discovery across research domains.

### Graph Structure Overview: UBKG, Petagraph, and the DDKG

The Data Distillery Knowledge Graph (DDKG) is implemented using a schema based on the NIH Unified Medical Language System (UMLS) implemented in the Unified Biomedical Knowledge Graph (UBKG), a scalable, ontology-aware property graph framework designed to support integration, traversal, and reasoning across heterogeneous biomedical data. The underlying model organizes knowledge as a directed graph of nodes (i.e. Concepts, Codes, Terms) and edges (semantic relationships), each annotated with controlled identifiers, source provenance, and ontology contexts.

Nodes in the schema are classified into core types:

● **Concept nodes** represent biomedical entities (e.g., genes, proteins, diseases, metabolites, anatomical regions) using identifiers from standard vocabularies (e.g., HGNC, MONDO, CHEBI).
● **Code nodes** are atomic identifiers (e.g., HGNC:11892, MONDO:0004975), and act as leaf nodes connected to Concept nodes.
● **Term nodes** represent lexical labels, synonyms, or text-derived phrases associated with Concept or Code nodes.

Each Concept node may be linked to one or more Code nodes (supporting identifier normalization), and to Term nodes that provide synonym, label, or lexical match support. This separation allows the graph to simultaneously support symbolic reasoning, lexical matching, and ontology alignment.

Edges in the graph are typed and directionally labeled to indicate relationship semantics (e.g., *expressed_in*, *part_of*, *regulates*, *treated_by*). Each edge is annotated with its source **`SAB`** (Source Abbreviation), such as GTEx, DrugCentral, or GlyGen, along with optional evidence codes or statistical weights.

The entire system is harmonized using ontologies drawn from over 180 curated biomedical vocabularies and terminologies, including UMLS (Unified Medical Language System), OBO Foundry ontologies, and CFDE-aligned data standards. Nodes and edges are semantically resolved through UMLS CUIs and graph-aligned identifiers, allowing concept equivalence and hierarchy-aware reasoning.

The DDKG extends the UBKG graph model by incorporating additional sources, including Common Fund datasets and CFDE-supplied annotations. These and other sources are incorporated into this core schema, enabling large-scale integration of omics data types (e.g., gene expression, chromatin loops, eQTLs, pathways, metabolites). Each dataset ingested into the DDKG is harmonized through this schema, ensuring consistency in concept typing, identifier resolution, and edge annotation.

This structured, modular approach enables the DDKG to support graph-native querying, shortest-path inference, and subgraph projection, while remaining extensible for downstream analytics and machine learning applications.

Development of the DDKG required a systematic approach to integrate datasets from multiple NIH Common Fund Data Coordinating Centers (DCCs) ^88^. The datasets varied in type, scale, and complexity, and were harmonized to standardized ontologies and incorporated into the graph to enable efficient querying and hypothesis generation. Building upon the Petagraph framework and the Unified Biomedical Knowledge Graph (UBKG), the DDKG employs a robust data integration process, schema customization, and curated knowledge ingestion to create a scalable and flexible knowledge graph platform. The next section outlines the datasets, data harmonization protocols, integration framework, and user interface development.

Versioning of the DDKG follows a date-based convention. Each release is distributed as a compressed archive named in the format *DataDistillery_dd_MMM_yyyy.zip*, where *dd_MMM_yyyy* corresponds to the date when the distribution was built and uploaded. The work presented in this paper is based on the DDKG January 2025 release (DataDistillery_03_JAN_2025.zip). This can be downloaded at the Data Distillery - Publication section within the download site at https://ubkg-downloads.xconsortia.org/ and will require registration and a free UMLS key to download at https://www.nlm.nih.gov/research/umls/index.html. Documentation is available in the NIH CFDE GitHub repository at https://github.com/nih-cfde/data-distillery where documentation for this release can be found under the DataDistillery03Jan2025 directory. Release notes accompany each distribution, and the complete archive of release notes is available in the ubkg-etl GitHub repository at https://github.com/x-atlas-consortia/ubkg-etl/tree/main/generation_framework/release%20notes/Data%20Distill ery.

For clarity, we indicate DDKG graph elements in bolded courier font in the rest of this report. For example, distinguishing graph elements such as **`HGNC`** nodes from the dataset source HGNC ^89^.

### 4.1 Common Fund Datasets Employed in the DDKG

The DDKG integrates a variety of datasets originating from different NIH Common Fund Data Coordinating Centers (DCCs) in addition to many other supporting datasets. A complete data dictionary for the Common Fund data sets can be found here: https://github.com/nih-cfde/data-distillery/blob/main/DataDistillery03Jan2025/DD_03Jan2025_userguide.md and review of the Data Distillery context in the UBKG can be found here: https://ubkg.docs.xconsortia.org/contexts/#data-distillery-context

Below we briefly summarize the Common Fund content in the DDKG. A full list and details can be found at the NIH-CFDE GitHub repository ^88,90^ as well as the UBKG Source Contexts documentation ^91^

● **4D Nucleome (4DN):** Chromatin loops from Hi-C experiments (https://commonfund.nih.gov/4Dnucleome) ^14^ from https://data.4dnucleome.org/
● **Extracellular RNA Communication Consortium (ERCC):** Loci where RNA binding proteins (RBPs) interact with extracellular RNA (exRNA) and the biofluids where these RBPs are predicted to be present. (https://commonfund.nih.gov/exrna)^15,16^ from https://exrna-atlas.org/
● **GlyGen:** Data from glycomics and proteomics databases (https://commonfund.nih.gov/Glycoscience) ^18^ from https://glygen.org/
● **Library of Integrated Network-based Signatures (LINCS):** Impact of drugs and small molecules on the expression of human genes in different cell lines (https://commonfund.nih.gov/lincs) ^23^ from (source websites)
● **Genotype-Tissue Expression (GTEx):** Bulk RNA-seq gene expression levels and eQTLs from GTEx version 8 (https://commonfund.nih.gov/GTEx) from https://www.gtexportal.org/home/index.html
● **Human BioMolecular Atlas Program (HuBMAP):** Cell-type-specific gene markers from single-cell data (https://commonfund.nih.gov/HuBMAP) from Azimuth https://azimuth.hubmapconsortium.org/
● **Illuminating the Druggable Genome (IDG):** Relationships between compounds, diseases, and molecular targets (https://commonfund.nih.gov/idg) ^21^ from https://druggablegenome.net/
● **Metabolomics Workbench (MW):** Relationships between metabolites, genes, cells, and diseases (https://commonfund.nih.gov/metabolomics) from https://www.metabolomicsworkbench.org/
● **MoTrPAC:** Omics data from studies on mechanisms of physical activity and health improvement, (https://commonfund.nih.gov/motrpac) from https://www.motrpac.org/
● **Kids First:** Phenotypes and high-impact variants per gene in pediatric cohorts (https://commonfund.nih.gov/KidsFirst) from https://portal.kidsfirstdrc.org

### 4.2 DDKG Schema Design and Adaptation

The DDKG schema builds upon the Petagraph KG, which integrates diverse biomolecular data using the Unified Biomedical Knowledge Graph (UBKG) annotation-rich infrastructure ^10^. Key design principles of the DDKG schema include:

1. **Extensibility**: The DDKG schema is inherently adaptable and extensible, supporting the integration of novel data types and ontologies from a wide array of biomedical sources. Users can utilize the DDKG as-is for querying integrated datasets or extend it by adding their own data. The schema supports seamless customization and expansion by leveraging UBKG ingestion protocols to incorporate new data, allowing DDKG to scale without disrupting existing data structures as new datasets and ontologies are added.
2. **Maintenance of Dataset Relationships**: A core feature of the DDKG schema is its ability to preserve and leverage relationships across datasets. This ensures researchers can perform cross-domain analyses without losing contextual integrity. For example:

1. ● **eQTL and Chromatin Interactions (example: Method 1)**: The schema connects eQTLs from GTEx with chromatin interaction data from 4DN, enabling integration with gene regulation.
2. ● **Drug-Target Pathways (example: Method 4)**: Drug-target relationships from IDG are linked with pathway-level annotations from LINCS, facilitating the interpretation of drug effects within broader biological pathways.
3. **Use of Standardized Ontologies**: By leveraging over 180 biomedical ontologies, including HPO and GENCODE, the schema ensures a consistent representation of biological entities. Ontology mapping anchors entities such as genes, proteins, and metabolites to standardized concepts, empowering cross-dataset queries and supporting compatibility with external knowledge graphs or ontologies.
4. **Graph Representation**: The schema represents biological entities as nodes, with edges denoting relationships such as “interacts with” or “expressed in.” Each relationship is expressed as a subject-predicate-object triplet, a model that enhances queryability and compatibility with graph-based machine learning techniques. This structured representation supports a wide range of analytical workflows, from simple queries to advanced graph-based computation.
5. **Scalability**: The DDKG schema is designed to scale dynamically, leveraging the UBKG’s automated pipelines to generate and harmonize nodes and edges efficiently. Its flexible architecture supports the inclusion of new datasets or relationship types without disrupting existing structures. This scalability ensures that the DDKG can grow alongside evolving research needs, supporting both immediate and future use cases.

### 4.3 DDKG Construction

The DDKG’s construction involves transforming datasets into graph-based data structures where nodes represent biological entities (genes, proteins, diseases, etc.) and edges represent relationships between these entities. We employ the UBKG ingestion protocols ^92^ that employ the PheKnowLator method^10,11^. The UBKG ingestion scripts extract biological relationships from disparate datasets and convert them into a format that can be represented within the graph. The UBKG scripts automate several stages of the DDKG construction:

#### 4.3.1 Ontologies and Standards

Data from contributing datasets is harmonized to ontologies approved by the CFDE Ontology Working Group. These include the Human Phenotype Ontology (HPO) and community standards like GENCODE. The UBKG ingestion scripts (**Section 4.3.4**) automate ontology mapping by assigning data identifiers with standardized ontology concepts in the graph. This process resolves data discrepancies, ensuring entities are uniformly represented across all sources. For example, chromosomal positions from multiple assemblies (e.g., GRCh37, GRCh38) are standardized to a unified coordinate system, while gene and protein identifiers are normalized to their canonical ontology terms (e.g., Ensembl, UniProt). This process ensures consistent entity representation across datasets (e.g., for genes, proteins, metabolites, compounds). It also ensures integration across diverse data types within a unified framework.

A key component of the biomolecular data harmonization process in the DDKG is to link genomic features’ chromosomal positions to the Homo Sapiens Chromosomal Location Ontology (HSCLO) ^29^, which was specifically developed to facilitate the integration of genomic data into biomedical knowledge graphs. HSCLO provides a structured representation of chromosomal locations, supporting integration and cross-querying at multiple data resolution levels. The HSCLO supports:

● **Hierarchical Resolution**: Nodes represent chromosomal locations at varying levels of detail (e.g., 1 Mb, 100 kb, 10 kb, and 1 kb), enabling efficient scaling and querying.
● **Cross-Dataset Integration**: Enables seamless linking of genomic features, such as chromatin loops from 4D Nucleome and gene annotations from GTEx or Kids First.
● **Ontology Standardization**: Harmonizes positional data to standardized genome assemblies (e.g., GRCh37 and GRCh38), ensuring compatibility across datasets.

HSCLO’s hierarchical relationships and structured node connections allow DDKG users to integrate genomic data effectively in their queries.

#### 4.3.2 Pre-Ingestion Protocols

To ensure the DDKG efficiently integrates data across diverse sources, we employ pre-ingestion data harmonization protocols. These protocols address data heterogeneity by aligning datasets with standard ontologies and formats, which supports FAIR principles, cross-dataset interoperability and consistent representation of biological entities. Before ingestion into the DDKG, datasets undergo:

1. **Data Cleaning**: Removing redundancies,ensuring data quality, and removing unnecessary data.
2. **Format Conversion**: Standardizing file types and formats to ensure compatibility with the ingestion pipeline.
3. **Schema Mapping**: Checking for alignment of data fields with the DDKG schema to preserve dataset relationships and facilitate querying.

#### 4.3.3 Data Integration Framework

The DDKG was built based on earlier work that integrated genomics data with the UBKG ^10^, incorporating data from the National Library of Medicine’s Unified Medical Language System (UMLS) and various external sources such as NCBO Bioportal and OBO Foundry. The UBKG Data Integration Framework provides several key components to the function of the DDKG:

1. **Source Context**: Collection of assertion sets identified by Source Abbreviations (**`SAB`**s).
2. **Source Framework**: Scripts that generate UMLS CSVs from UMLS data, integrating additional assertion data from external sources.
3. **Generation Framework**: This framework extends the UMLS context by incorporating additional assertions to create ontology CSVs, which are then imported into Neo4j databases.
4. **Safe API**: Provides access to the ontology knowledge graph database, abstracting common query types (**Section 2.4**)

#### 4.3.4 UBKG Ingestion Protocols

DDKG dataset ingestion is automated using UBKG scripts, enabling large-scale processing of datasets while maintaining biological context. These scripts also allow dynamic updates to integrate new datasets as they become available. The UBKG ingestion framework automates key steps in data harmonization and integration:

1. **Ontology-Based Mapping:** Uses predefined mappings from UBKG’s ontology libraries to ensure data compatibility and accuracy. The mapping process automatically aligns dataset entities with ontological terms. During ingestion, the UBKG scripts first map biological entities (such as genes, proteins, and diseases), to the graph’s corresponding Concept nodes provided by standard biomedical ontologies. This ensures that data from different sources are harmonized and connected through standard terminologies.
2. **Triples Generation:** Converts biological relationships into subject-predicate-object triplets for graph representation. The UBKG scripts generate the nodes (biological entities) and edges (relationships) that will form the structure of the DDKG. Each biological relationship is represented as a subject-predicate-object triplet, where the subject and object are biological entities (e.g., genes or diseases) and the predicate describes the type of relationship between them (e.g., "associated with" or "regulates"). These triplets are then translated into CSV files, which serve as the input for the Neo4j graph database.
3. **Data Transformation and Bulk Import:** The generated CSV files are tailored for bulk ingestion into the **Neo4j** database using the **Neo4j-admin import tool**. The UBKG scripts ensure that all datasets are formatted correctly, with each file conforming to Neo4j’s requirements for node and edge definitions. The transformation process integrates multiple datasets into the graph in a way that preserves the biological meaning and context of the data while maintaining a consistent schema across all sources.

By leveraging these automated processes, the DDKG can efficiently integrate diverse datasets into a unified graph structure while maintaining scalability and adaptability. Information on the UBKG and its ingestion protocols can be found at https://ubkg.docs.xconsortia.org/

#### 4.3.5 Manual Cross-Dataset Compatibility Checks

While our automated harmonization protocols ensure consistent formatting and ontology alignment, cross-dataset compatibility is manually assessed to maintain biological relevance and contextual accuracy. We manually verified the ability to do cross-dataset queries, for example, between eQTLs from GTEx and chromatin loop data from 4D Nucleome to investigate regulatory interactions, or disease pathway integrations between Metabolomics Workbench profiles and LINCS gene expression data.

### 4.4 Application Notes

#### 4.4.1 Method 1 Application: Querying Overlapping Chromosomal Features

**Query 1** is an example of a spatially resolved, graph-native strategy for investigating chromosomal feature co-localization. Typical methods for detecting feature overlaps is file-based parsing and interval joins, but here the DDKG encodes chromatin structures and eQTLs as semantically linked Concepts aligned to fixed-resolution genomic bins (**`HSCLO`** nodes, 1 kbp). Using the Cypher **`shortestPath()`** operator, the query efficiently identifies the span of genomic bins between **`4DNL`** loop anchors. By matching **`GTEXEQTL`**s to any bin intersecting that span, it reveals GTEx eQTL variants likely positioned within functionally constrained chromatin neighborhoods from 4DN data. This type of query on the DDKG enables high-resolution integration of chromosomal features without external annotation tools.

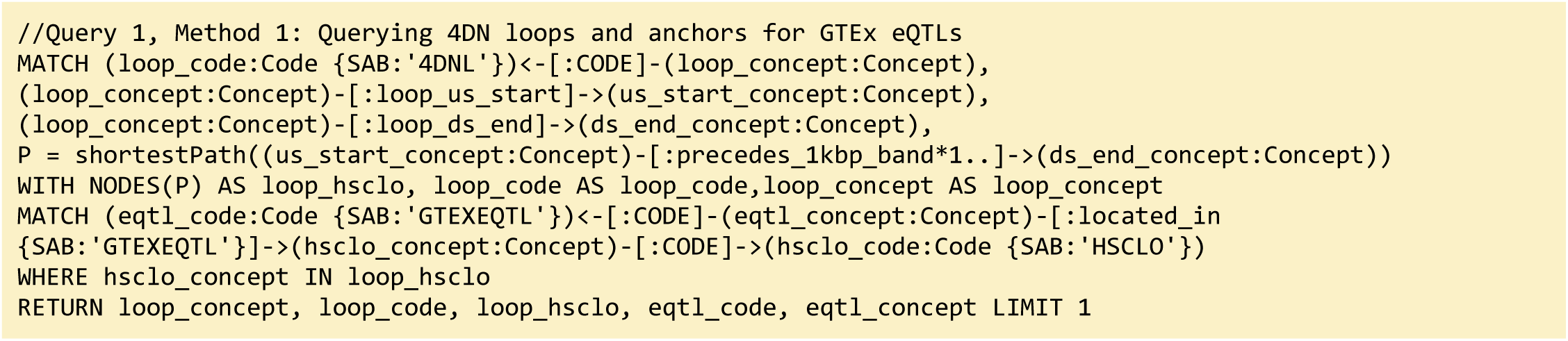

**Query 1 Description:** This Cypher query explores the relationships between a specific genomic Concept (**`4DNL`**) and eQTL (expression quantitative trait loci) data from the **`GTEXEQTL`** dataset. We start by identifying the **`loop_concept`** associated with the **`4DNL`** Code and its connections to two other Concepts: **`us_start_concept`** (the start of a loop) and **`ds_end_concept`** (the end of the loop). We then compute the shortest path between these two Concepts, capturing the set of relevant **`HSCLO`** Concepts (**`loop_hsclo`**) along the way. Next, we match **`eqtl_concept`** Concept nodes connected to the **`GTEXEQTL`** Code and filter for those linked to the variable **`hsclo_concept`** (high-scoring loci) that is part of the loop_hsclo set. Finally, we return results of variables **`loop_concept`**, **`loop_code`**, **`loop_hsclo`**, **`eqtl_code`**, and **`eqtl_concept`** to uncover any potential associations between the loop structure and eQTL data, limiting the output to one result.

Figure 2 in the main text presents an example output from **Query 1**, demonstrating an overlap between the upstream anchor start site of a representative **`4DN`** chromatin loop and a **`GTEx`** eQTL Concept node. This result highlights how **`HSCLO`** Concept nodes at 1 kbp resolution can be utilized for integrating diverse genomic features by enabling the discovery of biologically meaningful associations across independent datasets. A deeper analysis of large-scale outputs from this query may provide insights into genomic regions where some co-occurrence events are highly prevalent, potentially identifying regulatory mechanisms underlying gene expression. The concurrent analysis of eQTL data and DNA loop information enables the identification of eQTLs located within regulatory elements that physically interact with their cognate target genes, establishing a more direct link between genetic variation and its impact on transcriptional output. This integrated approach also facilitates the resolution of causal variants from those in linkage disequilibrium and the precise identification of target genes influenced by eQTLs, especially in cases of significant linear genomic separation. Furthermore, examining eQTLs within the context of DNA loop structures elucidates the underlying molecular mechanisms of gene regulation, including the impact of genetic variation on transcription factor binding and enhancer-promoter interactions. Other studies underscore the significance of higher-order chromatin organization in mediating distal gene regulation by facilitating the spatial co-localization of eQTLs and their target genes.

##### Query 1 Semantic Path

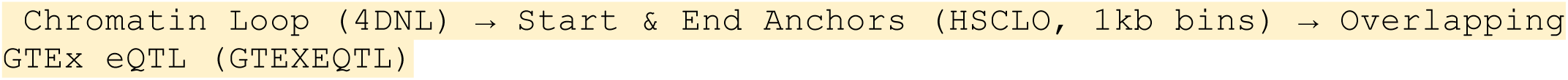

This path represents the traversal from a chromatin loop, through its upstream and downstream genomic anchors (at 1 kbp resolution), to any eQTLs located within those anchors. It illustrates how the DDKG enables proximity-based feature discovery by semantically connecting structural and regulatory genomic elements.

The results from Query 1 can be found in **Supplementary File 1**.

#### 4.4.2 Method 2 Application: Identifying Genes, Diseases, and Biomarkers

**Query 2** demonstrates a multi-hop graph traversal connecting disease phenotypes to molecular biomarkers in accessible biofluids using ontology-based relationships. While typical methods usually query isolated data tables, the Cypher query identifies genes associated with a specified Human Phenotype Ontology (HPO) Term, traces their exRNA expression loci from ENCODE annotations, and resolves their potential detectability through known RNA-binding proteins (RBPs) present in specific UBERON-anatomy-coded biofluids. The use of semantically labeled edges such as **`overlaps, molecularly_interacts_with, and predicted_in`** provides constraints for the search. By linking **`ERCCRBP`** binding predictions to tissue localization in **`UBERON`**, this graph-native approach supports construction of targeted, interpretable workflows for non-invasive biomarker discovery. It results in a path from phenotype to gene to exRNA-binding RBP to biofluid, all by using ontologies in the query logic without external preprocessing. This approach generalizes to any disease phenotype with linked data and supports hypothesis generation for any biomarker project.

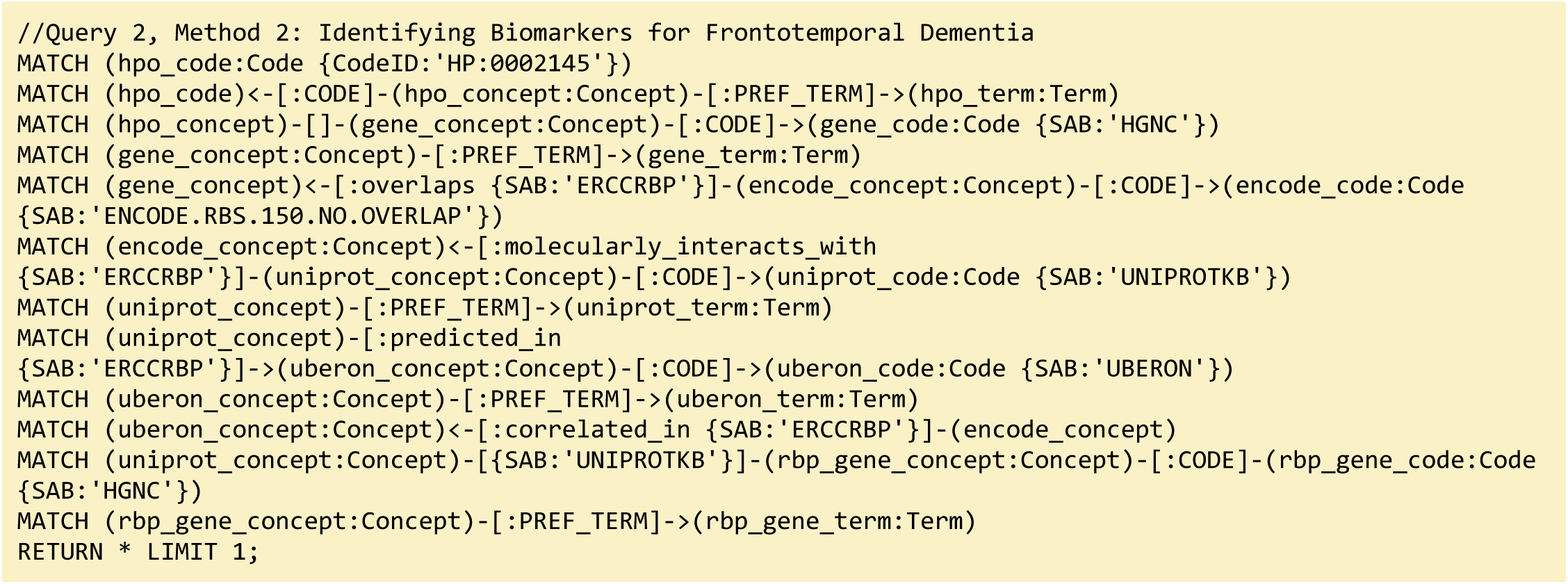

**Query 2 Description:** The query starts by identifying the **`HPO`** Code for Frontotemporal Dementia with **`CodeID`** of **`HP:0002145`**, which is connected to its appropriate **`hpo_concept`** that is linked to **`hpo_term`** via the **`PREF_TERM`** relationship. The **`hpo_concept`** node is then connected to a **`gene_concept`** node (with **`gene_term`** and **`gene_code`** for each gene), linking disease to gene. Since we are interested in finding the loci within those genes where exRNA expression is detected, the query connects **`gene_concept`** with **`encode_concept`** via **`overlaps`** relationship. In addition, we identify **`encode_code`** and confirm that we selected appropriate nodes by checking that their **`SAB`** value is **`ENCODE.RBS.150.NO.OVERLAP`**. In order to detect the biofluids where exRNA loci expression was detected we have connected the nodes **`uberon_concept`** and **`encode_concept`** using the **`correlated_in`** relationship. The variable **`uberon_concept`** is connected to **`uberon_term`** and **`uberon_code`**, which checks that the **`SAB`** value for Code nodes is **`UBERON`**. To detect the RBPs predicted to be present in that biofluid we have to find **`uniprot_concept`** connected to **`encode_concept`** via **`molecularly_interacts_with`** relationship, that are at the same time connected to **`uberon_concept`** via **`predicted_in relationship`**. The **`uniprot_concept`** Concept nodes are connected to **`uniprot_term`** and **`uniprot_code`** to check that Code nodes **`SAB`** values are **`UNIPROTKB`**. As a final step in the query, we select RBP genes as **`rbp_gene_concept`** that is connected to **`uniprot_concept`** and further identify **`rbp_gene_term`** and **`rbp_gene_code`**, where Code nodes have **`SAB`** value **`HGNC`**. We limit returns to one set of data for better graphical representation of the results. This query can be changed to search for results for any phenotype or disease by changing just the **`CodeID`** value for the **`hpo_code`** at the beginning of the query.

##### Query 2 Semantic Path

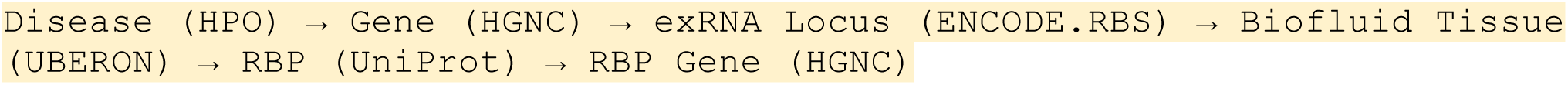

**Query 2’s** path traces from a disease phenotype to its associated genes, identifies where their exRNA products are observed, detects the RNA-binding proteins predicted to bind these exRNAs, and then links those proteins to the tissues or biofluids in which they are predicted to be present. It also resolves the coding genes for those RBPs, providing a full biomarker hypothesis path.

**Query 3** is an example of how the DDKG can be used to evaluate molecular consequences of pharmacologic perturbation by tracing a drug-gene-exRNA-RBP-biofluid path through the graph. It begins with a specific **`PUBCHEM`** compound, here Astemizole, and identifies genes upregulated in **`LINCS`** perturbation datasets. The gene nodes are linked to genomic loci associated with extracellular RNA records and intersecting RNA-binding protein (**`ERCCRBP`**) targets. Rather than manually intersecting genomic intervals or transcriptomic signatures, the query uses the DDKG’s curated **`overlaps, molecularly_interacts_with,`**and **`predicted_in`** relationships to traverse from drug perturbation to potential exRNA biomarkers. It further filters RBPs based on their predicted localization to specific UBERON-encoded anatomical fluids, enabling a direct link from compound activity to biofluid-accessible targets. The query supports inference of molecular mechanisms and facilitates the design of liquid biopsy strategies to monitor treatment response. This combines functional genomics, pharmacology, and tissue-specificity in a single multi-hop query, demonstrating the power of the DDKG for systems pharmacology and translational discovery.

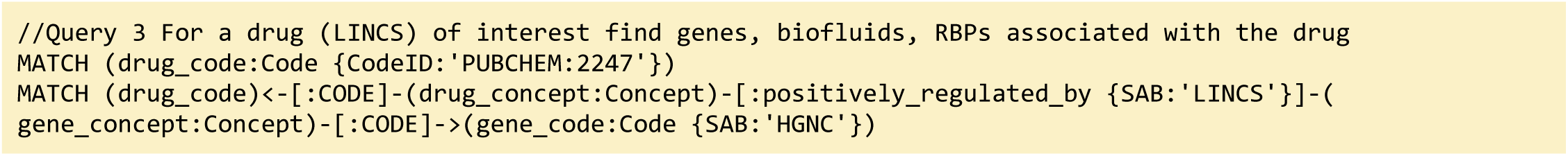

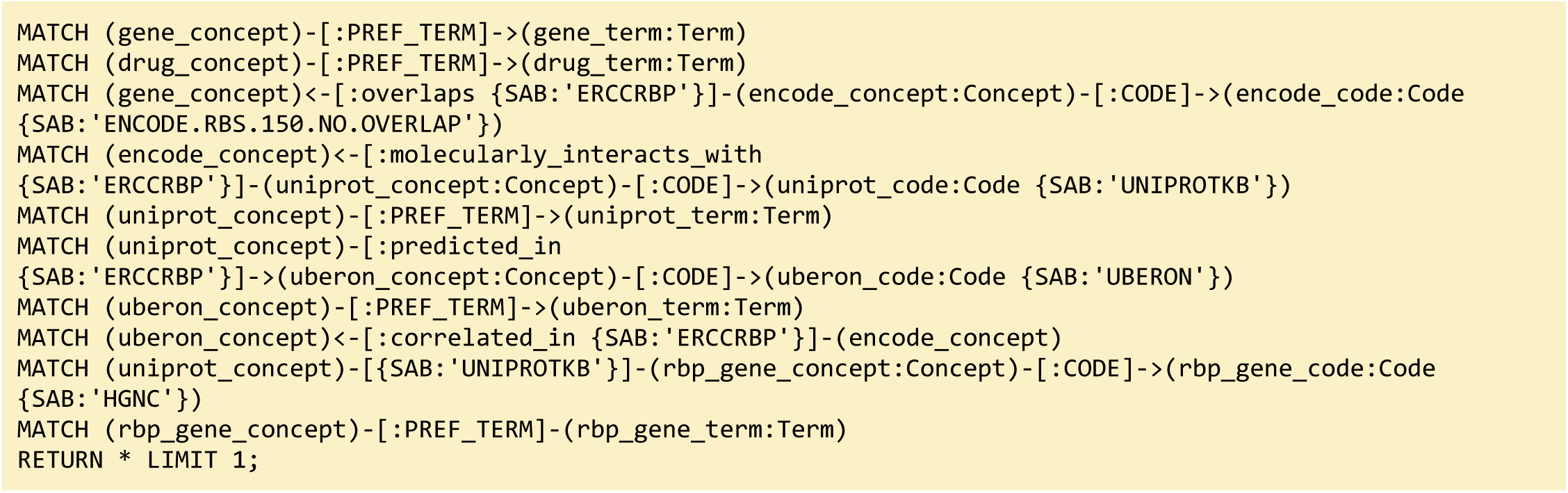

**Query 3 Description:** The query starts by identifying the exact **`drug_code`** for the drug of interest, in this case **`Astemizole`**, with **`CodeID PUBCHEM:2247`**, which is connected to its appropriate **`drug_concept`** that is linked to **`drug_term`** via the **`PREF_TERM`** relationship. Next we need to find genes associated with that drug by finding **`gene_concept`** connected to **`drug_concept`** via **`positively_regulated_by`** relationship. The **`gene_concept`** is connected to **`gene_term`** and **`gene_code`**, where Code nodes have **`SAB`** value **`HGNC`**. As in the previous query, we are seeking loci within those genes where exRNA expression is detected by connecting gene_concept with encode_concept via **`overlaps`** relationship, and finding **`encode_code`** where the **`SAB`** value is **`ENCODE.RBS.150.NO.OVERLAP`**. To identify RBPs, we identify **`uniprot_concept`** connected to encode_concept via **`molecularly_interacts_with`** relationship. The **`Uniprot_concept`** relates to **`uniprot_term`** and **`uniprot_code`**, confirming that Code nodes **`SAB`** values are **`UNIPROTKB`**. To identify biofluids in which these RBPs are predicted to be present, we query **`uberon_concept`** connected to **`uniprot_concept`** via **`predicted_in`** relationship. The **`uberon_concept`** nodes are connected to their **`uberon_term`** and **`uberon_code`** nodes, confirming that the **`SAB`** value for Code nodes is **`UBERON`**. In order to select only the biofluids where exRNA loci expression is detected we have to find **`uberon_concept`** connected to **`encode_concept`** using **`correlated_in`** relationships. Finally, we select RBP genes as **`rbp_gene_concept`**, connected to **`uniprot_concept`** and find their **`rbp_gene_term`** and **`rbp_gene_code`**, where Code nodes have **`SAB`** value of **`HGNC`**. Results are limited to just one set of data for visualization purposes. This query can be changed to search for results for any other drug by changing just the **`CodeID`** value for **`drug_code`** at the beginning of the query (Figure 8).

##### Query 3 Semantic Path

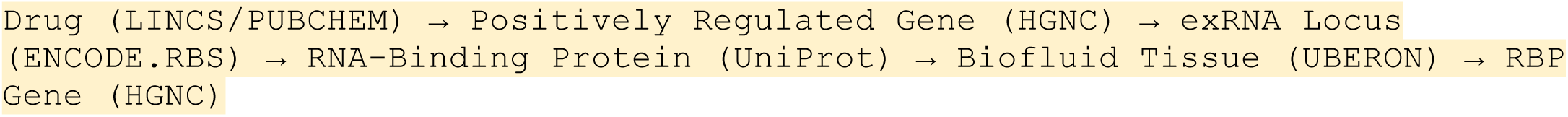

This path traces from a drug to the genes it positively regulates, identifies where those genes produce exRNAs, detects the RBPs that bind those exRNA loci, then finds the tissues or biofluids where those RBPs are predicted to be present. It ends by resolving the coding genes for the RBPs themselves, forming a full loop from perturbation to detection strategy.

#### 4.4.3 Method 3 Application: Tissue Specific Expression of Glycosylation Enzymes

**Query 4** focuses on linking glycosylation enzymes with GTEx expression and tissue ontologies. This approach avoids manual identifier conversions and provides an end-to-end semantic mapping in a single graph-native query. This results in a rapid, scalable screening of glycoenzyme expression across tissues, facilitating comparative and translational glycomics analyses.

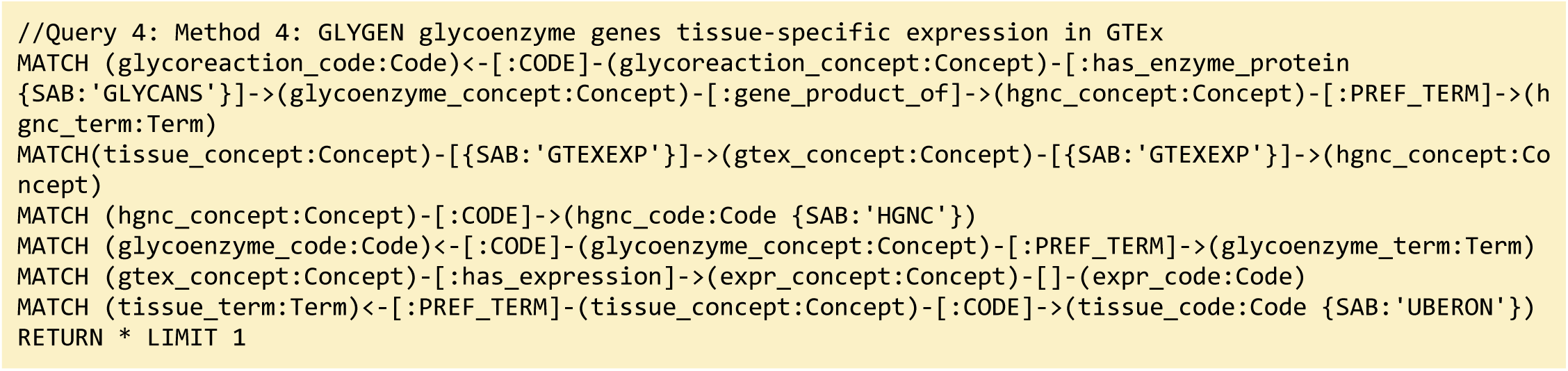

**Query 4 Description:** This query identifies relationships between glycoreactions, enzymes, gene products, and tissue expression data. The following Cypher query is designed to explore connections between glycoreactions, enzymes, tissues, and gene expression data, specifically linking these entities through various Concepts, Codes, and Terms. The query starts by identifying a **`glycoreaction_code`**, which is associated with a **`glycoreaction_concept`**. This Concept is connected to a **`glycoenzyme_concept`** via the **`has_enzyme_protein`** relationship, which is further linked to a **`hgnc_concept`** (a gene symbol) through a **`gene_product_of`** relationship. The **`hgnc_concept`** is then connected to a **`hgnc_term`** via the **`PREF_TERM`** relationship. We also want to explore expression data from the **`GTEXEXP`** dataset, where a **`tissue_concept`** is linked to a **`gtex_concept`** (representing gene expression data in tissues). The **`gtex_concept`** is also connected to the **`hgnc_concept`**, showing the relationship between gene expression and specific genes. We also identify the **`hgnc_code`** and **`glycoenzyme_code`** in order to return further details on gene and enzyme annotations. Additionally, the **`glycoenzyme_concept`** is connected to a **`glycoenzyme_term`**, and the **`gtex_concept`** is related to **`expr_concept,`** representing expression data, with connections to relevant Codes for expression and tissue data (e.g., **`UBERON`**). In short, we aim to uncover the relationships between glycoreactions, glycoenzymes, gene products, tissue-specific expression data, and gene annotations, in order to map how glycosylation and gene expression are linked across various biological contexts.

##### Query 4 Semantic Path

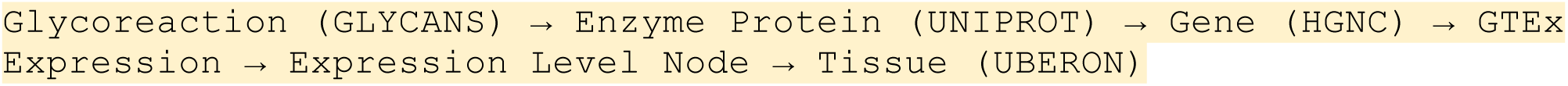

This path starts at a curated glycoreaction, traces through its catalyzing enzyme and associated gene product, and connects to expression levels recorded in the **`GTEx`** dataset. Those expression nodes are mapped to anatomical tissues via **`UBERON`** Terms. The semantic design allows users to analyze tissue-specific activity of enzyme-coding genes in the context of glycosylation pathways.

#### 4.4.4. Method 4 Application: Identifying Genes and Associated Drug Targets

In this use case, we perform three DDKG searches against lipoxygenase genes and asthma-associated drug targets. These queries use Cypher’s pattern-matching and list expansion capabilities to traverse multiple ontological and experimental relationships within the DDKG. In **Query 5**, we anchor the traversal to a phenotype label (e.g., ’Asthma’) using a text match on **`HP`** Term names rather than hard-coded concept identifiers to support flexibility and readability. The **`WITH`** clause allows dynamic injection of disease names as variables, enabling easy reuse across different disease terms. The query then exploits bi-directional traversal between Concept nodes and Code/Term nodes to identify **`HGNC`** coded genes associated with the phenotype and their perturbation links to **`PUBCHEM`** compounds through **`LINCS`** relationships. By limiting the traversal to specific source annotations using **`{SAB:’LINCS’}`**, it ensures that only curated experimental associations are returned. The multi-hop path leverages semantic edges to extract drug–gene–disease connections efficiently without intermediate steps.

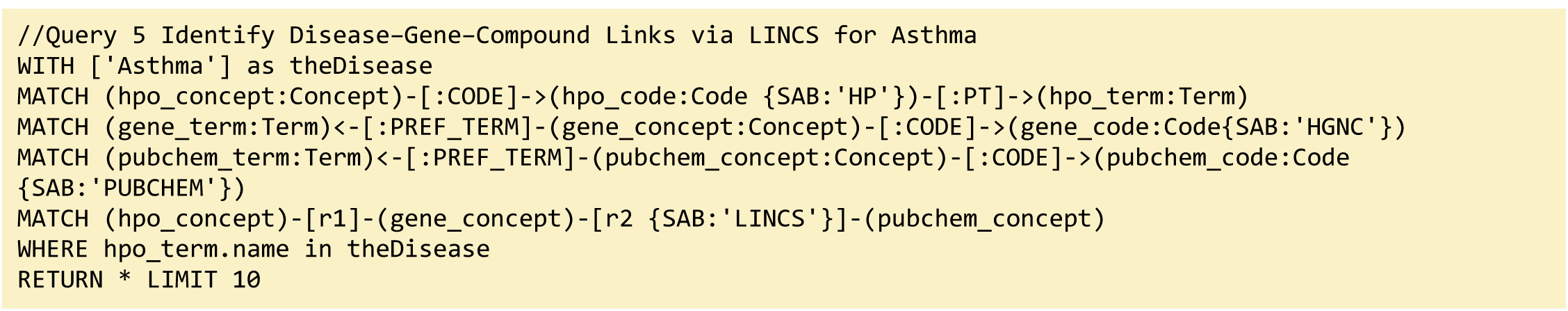

**Query 5 Description:** Combining data from IDG-DrugCentral and LINCS, this Cypher query on the DDKG seeks genes and Pubchem compounds associated with the disease “Asthma” (Figure 9). It then traverses from disease-linked HPO Concepts to associated genes (**`HGNC`**nodes), and from those genes to compounds (**PUBCHEM**-coded nodes) through **`LINCS`** relationships. This captures compounds that up- or down-regulate asthma-associated genes. The output includes all matching paths between asthma, lipoxygenase family genes (e.g., ALOX5), and relevant compounds that are known or potential inhibitors. This multi-hop traversal highlights the DDKG’s capacity to link across multiple datasets, in this case, disease phenotypes to pharmacological agents via transcriptomic perturbation data.

##### Query 5 Semantic Path

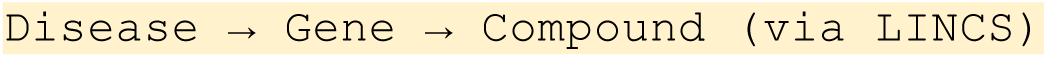

This semantic path begins with a disease phenotype, specified by matching text to a Human Phenotype Ontology (HPO) Term, and identifies genes associated with that phenotype in the DDKG. These disease-associated genes are then linked to compounds through transcriptomic perturbation data from LINCS. This multi-hop traversal—from phenotype to gene to compound—demonstrates the DDKG’s ability to integrate disease ontologies with pharmacogenomic annotations. It enables rapid identification of compounds that may up- or down-regulate disease-relevant genes, streamlining the discovery of candidate therapeutics or repurposing opportunities.

The central node in Figure 9, “893415,” is a gene named ALOX5 which is known to be involved with inflammation^93^. The result from a DDKG query shows that ALOX5 is negatively regulated by 39 compounds and positively regulated by 4 compounds.

The human genome has six functional lipoxygenase genes (ALOX15, ALOX15B, ALOX12, ALOX12B, ALOXE3, and ALOX5). **Query 6** searches for lipoxygenase enzymes within the DDKG containing the string “ALOX” and returns a sub-graph of enzymes and compounds associated via **`bioactivity`**.

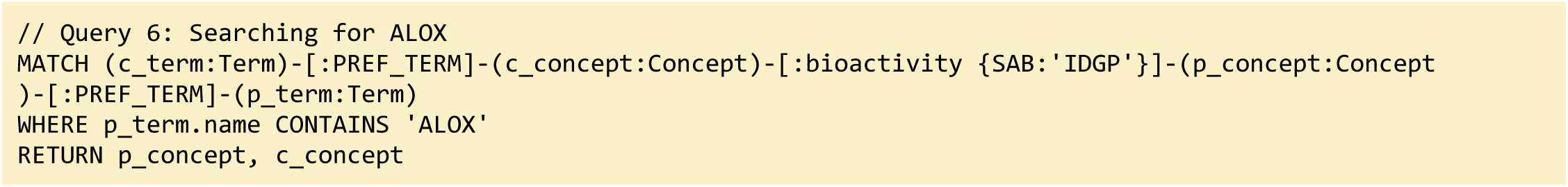

**Query 6 Description:** This query retrieves all bioactivity relationships from the **`IDG`**–DrugCentral dataset in the DDKG where the protein (gene) target contains the string “ALOX,” representing members of the lipoxygenase enzyme family. It matches compound Concepts connected via **`bioactivity`** edges to genes such as ALOX5 and ALOX15B. The result returns all compound–gene pairs with inferred activity relationships, supporting hypothesis generation for drug repurposing or mechanistic studies involving inflammation-linked pathways.

##### Query 6 Semantic Path

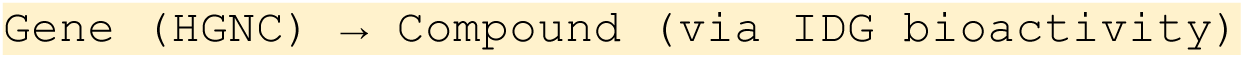

The **Query 6** semantic path traces from genes to bioactive compounds using direct bioactivity assertions from IDG-DrugCentral. The query filters for lipoxygenase family members by matching gene names containing “ALOX,” then retrieves all PubChem compounds with documented activity against these genes (Figure 10). By structuring genes and compounds as connected nodes within the DDKG and indexing their relationships through curated assertions, this query supports fast retrieval of compound–gene relationships for therapeutic targeting or chemical biology investigations. It highlights how the DDKG supports focused pharmacologic queries with high specificity and coverage.

In order to investigate the unique tissue types associated with the HGNC term “*ALOX5*”, we devise a query that combines data from both the IDG and the GTEx DCCs:

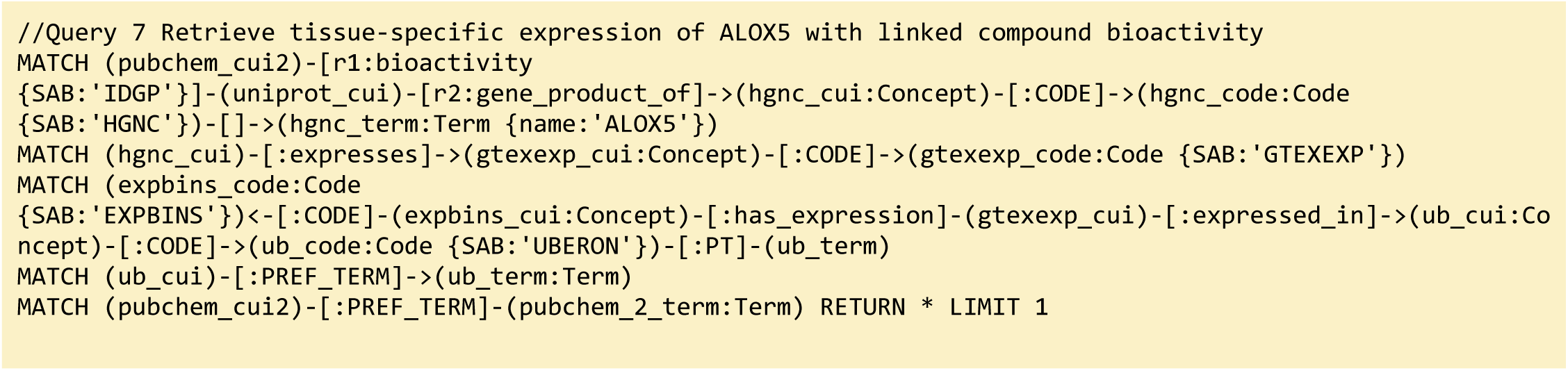

**Query 7 Description**: This query traces from the ALOX5 gene to known bioactive **`PUBCHEM`** compounds through **`UNIPROT`** gene products, and then to GTEx expression bins, which indicate approximate expression levels in different **`UBERON`** tissues. **`GTEXEXP TPM`** values are binned into discrete ranges (stored as **`EXPBINS`** nodes) to support faster and more interpretable semantic queries. By combining molecular, chemical, and anatomical contexts, this query enables tissue-aware exploration of drug targets and associated compounds in a single step (Figure 11).

##### Query 7 Semantic Path

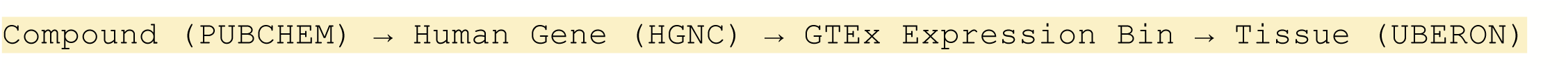

This approach facilitates queries such as identifying tissues where ALOX5 is highly expressed and cross-referencing them with known compound activity, supporting applications in drug repurposing, biomarker localization, and safety screening.

We find that there are several compounds associated with two different human ALOX enzymes. Specifically, ten compounds are associated with both ALOX15 and ALOX12, and one compound is associated with both ALOX15 and ALOX15B. One of the compounds associated with ALOX15B (PUBCHEM:1778842) is 3-[(4-Methylphenyl)methylsulfanyl]-1-phenyl-1,2,4-triazole (also known as “MLS000536924”) and was identified using high throughput screening as a competitive inhibitor (ki = 2.5+/-0.5 uM) of human epithelial 15-lipoxygenase-2 (ALOX15B) ^43^.

Running **Query 7** specifically for ALOX5 returns GTEx-derived expression bins linking ALOX5 to breast epithelium, Peyer’s patch, and anterior lingual gland—findings that are consistent with the literature on human ALOX5 expression. These graph-based results validate the integration of expression and compound annotations in the DDKG and support tissue-specific drug prioritization.

#### 4.4.5 Method 5 Application: Metabolites-related Studies

The MW schema enables the exploration of metabolites in relation to a condition or phenotype (disease-metabolite), sample source (cell-metabolite), gene (gene-metabolite), and vice versa. Further, the metadata for condition or phenotype, sample source, and gene can be used as anchors to explore and combine resources from other DCCs and gain insights beyond metabolomics. For instance, IDG resources can identify potential targets (genes or metabolites) and small molecules against those targets; LINCS resources can be used to predict gene expression signatures for the small molecules; exRNA for potential miRNA biomarkers; and so on. Encoding of nodes and their relationships with controlled terminology drives this harmonization and integration of disparate data using common metadata elements across CFDE, thus enhancing the value of common fund datasets. For example, for a disease, find drugs that may treat that disease (IDG), then find genes affected by that drug, and then find the metabolites related to those genes. In view of this, understanding how a compound influences biological systems, from its bioactivity on proteins to its effects on genes, metabolites, and conditions in specific tissues, is of great importance. By tracing the relationships between compounds, proteins, genes, metabolites, and conditions, researchers can gain insights into the underlying molecular mechanisms of drug action, disease pathways, and potential therapeutic targets. This type of information is crucial for identifying biomarkers, understanding metabolic networks, and exploring how metabolites are produced in tissues under different conditions, which could inform the development of new drugs or treatments. Additionally, linking standardized preferred terms to each Concept ensures that the data is consistent and interoperable, making it easier to integrate with other biological datasets or databases, ultimately advancing our understanding of complex biological systems. (Figure 4)

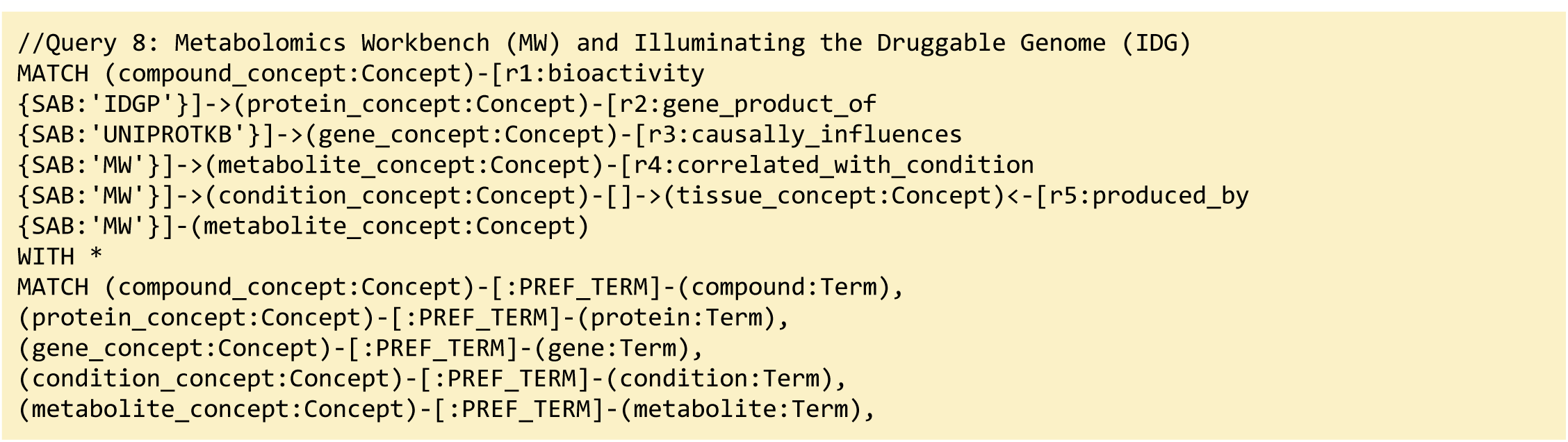

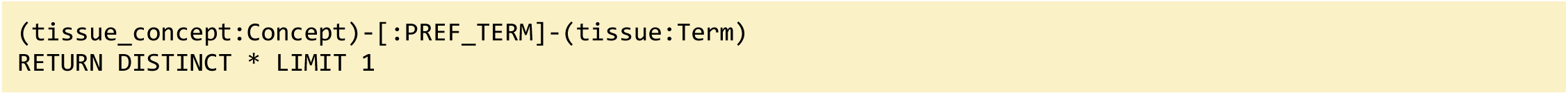

**Query 8 Description:** With this query, we are aiming to trace a specific biological pathway in the graph, starting from a compound with bioactivity, and following it through several key biological concepts: proteins, genes, metabolites, conditions, and tissues. Specifically, we seek to identify a compound’s bioactivity towards a protein (**`SAB`**: **`IDGP`**), link the protein to its corresponding gene (**`SAB`**: **`UNIPROTKB`**), and then track how the gene causally influences a metabolite (**`SAB`**: **`MW`**). We also want to identify conditions correlated with this metabolite and see how these conditions are associated with tissues that produce the metabolite. Additionally, we retrieve the preferred Terms (standardized names or identifiers) for each of these Concepts (compound, protein, gene, metabolite, condition, and tissue) to ensure consistency and clarity in our results.

##### Query 8 Semantic Path

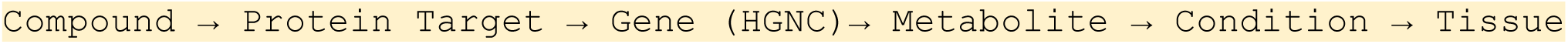

This semantic path traces the cascade of molecular and phenotypic associations from a pharmacological compound to its physiological context. The query begins with a compound known to exhibit bioactivity (from **`IDG`**–DrugCentral), identifies the protein target of that compound (via **`UNIPROT`**), and links it to the encoding gene. It then follows a causal influence relationship from the gene to a specific metabolite, using curated Metabolomics Workbench (**`MW`**) assertions. That metabolite is connected to disease or experimental conditions through **`MW`**’s **`correlated_with_condition`** relationship and ultimately mapped to tissues where the metabolite is produced. By chaining together bioactivity, gene regulation, metabolite dynamics, and tissue-specific production, this query enables multiscale exploration of drug mechanisms, metabolic signatures, and condition-specific biomarker identification. It demonstrates how the DDKG supports long-range biological inference by integrating chemically anchored relationships with gene–metabolite–phenotype axes in a unified graph.

#### 4.4.6 Method 6 Application: Tracing Expression Perturbations to Human Disease Phenotypes through Multi-Omic and Cross-Species Integration

Integrating multi-omic and cross-species data provides a comprehensive approach to understanding how experimental perturbations translate to human disease phenotypes. By tracing molecular signals from preclinical models (e.g., MoTrPAC rat data) through orthologous human genes, tissue-specific regulatory eQTLs (GTEx eQTLs), researchers can identify causal pathways linking genes to phenotypes and environmental or pharmacological interventions. This systems-level integration reflects strategies used in translational bioinformatics to prioritize mechanistic hypotheses and therapeutic targets, especially in complex diseases where single-omic data is insufficient. This integrated approach supports the discovery of biologically plausible links between experimental findings and clinical relevance, which is critical for translating omics data into actionable insights.

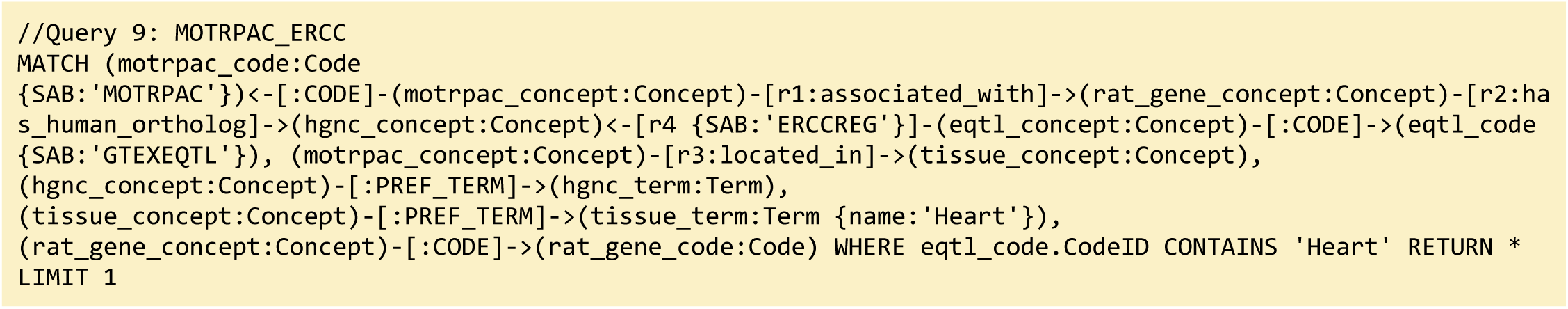

**Query 9 Description:** In this query, we aim to identify biologically meaningful links between **`MOTRPAC`**-coded concepts and human genes regulated by heart-specific eQTLs, thereby integrating rat preclinical data with human gene regulatory insights in a tissue-specific manner. The query begins by selecting Code nodes from the **`MOTRPAC`** source **`(motrpac_code:Code`** {**`SAB:’MOTRPAC’})`**, which are connected via the **`CODE`** relationship to broader biological Concept nodes (**`motrpac_concept`**). These Concepts are then traced through the **`associated_with`** relationship to corresponding rat gene Concepts (**`rat_gene_concept`**), signifying functional or experimental associations in the **`MOTRPAC`** dataset. From there, the query leverages the **`has_human_ortholog`** relationship to map these rat genes to their human orthologs (**`hgnc_concept`**). To connect these human genes to expression regulation data, the query uses the **`associated_with`** (specifically **`r4`** with {**`SAB:’ERCCREG’`**}) relationship to link the human gene Concepts to eQTL Concepts (**`eqtl_concept`**) curated in the **`ERCCREG`** resource. These eQTLs are then connected via the **`CODE`** relationship to GTEx eQTL Code nodes (**`eqtl_code`**) where the **`CodeID` i**ncludes "**Heart**", filtering for heart-specific regulatory variants. The original **`MOTRPAC`** Concepts are also filtered through the **`located_in relationship`** (**`r3`**) to a **`tissue_concept`** that is labeled with a **`PREF_TERM`** node (**`tissue_term`**) explicitly named "**Heart**", ensuring that only heart-related **`MOTRPAC`** findings are considered. Additionally, the human gene Concept is labeled with its preferred human-readable name through the **`PREF_TERM`** relationship to a Term node (**`hgnc_term`**), and the rat gene is similarly annotated via the **`CODE`** relationship to a **`rat_gene_code`**. Altogether, the query spans multiple ontologies and species, using well-defined relationship type **`associated_with`**, **`has_human_ortholog`**, **`located_in`**, to build a cross-species, heart-specific network. This enables identification of human genes regulated by heart eQTLs that are orthologous to rat genes functionally linked to heart-active molecular signatures from the **`MOTRPAC`** study, offering insights into conserved mechanisms of gene regulation and heart physiology.

##### Query 9 Semantic Path

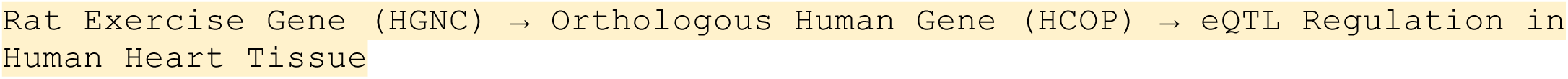

This semantic path begins with a MoTrPAC-identified exercise-responsive gene in rats, identified by the **`associated_with`** relationship from a **`MoTrPAC`** Concept node. The query follows a **`has_human_ortholog`** relationship to its corresponding human gene (**`HGNC`**), establishing a cross-species connection. From the human gene node, the path then traverses to GTEx-derived eQTLs that regulate gene expression in human heart tissue, using the **`ERCCREG`** relationship to link to eQTL Concept nodes, which are filtered for tissue specificity. This entire traversal is constrained to rat genes located in heart tissue and their human orthologs that are also eQTL-regulated in human heart samples. This multi-omic, cross-species path links experimental exercise biology in rodents to regulatory mechanisms in human tissues, offering a route for identifying conserved molecular pathways involved in cardiovascular adaptation. The query exemplifies how the DDKG enables mechanistic inference by bridging preclinical data, tissue-specific regulation, and orthology-based human relevance within a single semantic query framework.

#### 4.4.7 Method 7 Application: Genetic variants versus heart defect-linked genes

This use case cross-species integration of phenotypic and genetic data by linking mouse phenotype terms from the Mouse Phenotype Ontology (MP) to human genetic and clinical datasets through a multi-step ontological mapping. By first aggregating MP Concepts hierarchically related through “isa” relationships, the query captures a broad set of phenotypic traits associated with mouse genes via curated gene-phenotype annotations from MGI (Mouse Genome Informatics). These mouse genes are then connected to their human orthologs using high-confidence one-to-one mappings from the HCOP (HGNC Comparison of Orthology Predictions) database, ensuring evolutionary and functional relevance ^48^. The human orthologs are subsequently linked to Kids First entries and patient cohorts, leveraging the KF connectivities, to associate genetic variants with specific clinical groupings. This integrative approach supports the translational inference of human disease phenotypes from experimentally validated mouse models, aligning with established frameworks for cross-species data integration in functional genomics and precision medicine ^94,95^

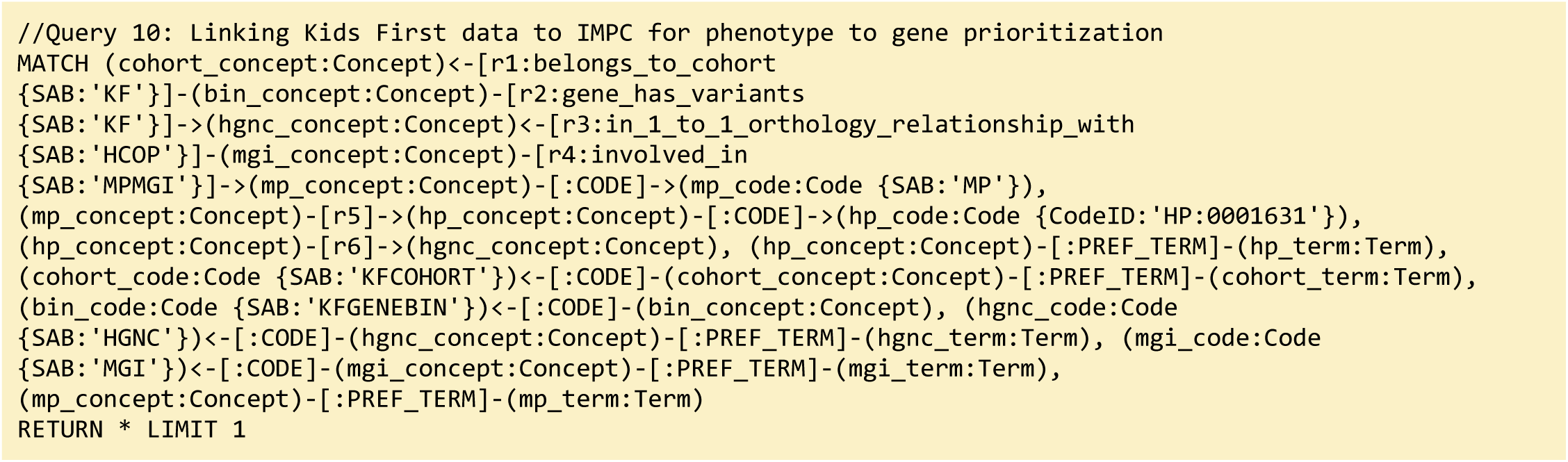

**Query 10 Description:** We aim to identify genes with rare variants in pediatric cohorts from the Kids First (**`KF`**) genomics program that may be implicated in cardiac abnormalities by leveraging cross-species orthology and phenotype ontologies. The query starts by retrieving gene variant bins (**`bin_concept`**) associated with KF cohorts (**`cohort_concept`**) using the **`belongs_to_cohort`** relationship, then links these bins to human genes (**`hgnc_concept`**) via the **`gene_has_variants`** relationship—indicating that these genes harbor variants in the cohort. Each human gene is mapped to its mouse ortholog (**`mgi_concept`**) through a high-confidence 1:1 orthology link (**`in_1_to_1_orthology_relationship_with`**, **`SAB: HCOP`**). These mouse genes are then connected to mammalian phenotypes (mp_concept) using the **`involved_in`** relationship (**`SAB: MPMGI`**), and the phenotype is anchored to formal Mammalian Phenotype Ontology Code nodes (**`mp_code`**). These mouse phenotypes are further mapped to human phenotypes (**`hp_concept`**), specifically filtered to include only those matching the Human Phenotype Ontology Term **`HP:0001631`** (Atrial Septal Defects), using the **`CODE`** relationship and supported by intermediate ontology links. A reverse link from the **`HP`** term back to the original human gene (**`hgnc_concept`**) helps reinforce cross-species relevance.

Preferred terms (**`PREF_TERM`**) and Codes (**`CODE`**) are retrieved throughout for cohort, bin, human gene, mouse gene, **`MP`**, and **`HP`** Concepts to support human-readable output. Through this multi-step ontology integration, the query highlights human genes carrying variants in rare disease cohorts that are supported by mouse model evidence and human phenotype mapping to be potentially relevant to congenital heart disease (Figure 13).

The full gene list can be found in **Supplementary File 2**.

##### Query 10 Semantic Path

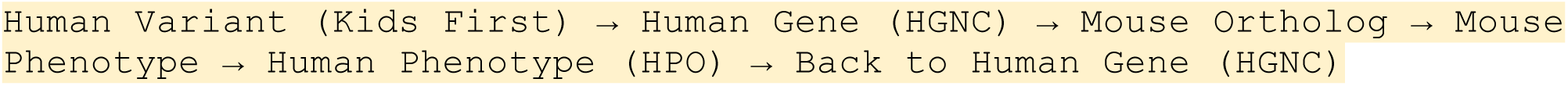

This multi-hop semantic traversal begins by identifying variant bins (**`bin_concept`**) associated with specific pediatric cohorts in the Kids First (KF) genomics program using the **`belongs_to_cohort`** `r`elationship. From each bin, it follows **`gene_has_variants`** edges to identify human genes (HGNC-coded). These human genes are linked to their murine orthologs using the **`in_1_to_1_orthology_relationship_with`** relationship from the HCOP database, ensuring high-confidence cross-species mapping. Mouse genes (MGI-coded) are then connected to Mammalian Phenotype Ontology (**`MP`**) Concepts using the **`involved_in`** predicate based on curated mouse knockout data from MGI/IMPC.

Each MP Concept is semantically bridged to a corresponding Human Phenotype Ontology (HPO) Concept using ontology-aligned cross-mappings. This path is filtered to retain only those linked to the HPO Code HP:0001631, corresponding to Atrial Septal Defect (ASD). A reverse mapping from the HPO Concept back to the original human gene reinforces the connection between patient variants and model-organism-supported phenotypic outcomes. By combining genetic data from human cohorts with phenotype annotations from mouse models, this path supports translational gene discovery for congenital heart defects via integrated ontology-based reasoning.

##### Validation with external enrichment analysis

To externally validate 27 genes identified from DDKG-based overlap between CHD cohort variants and IMPC phenotype-matched orthologs, we used those genes to perform enrichment analysis using the STRING (v12.0) database at https://string-db.org. STRING applies a one-sided hypergeometric over-representation test to assess whether functional categories occur more frequently in the input gene list than expected by chance, with multiple-testing correction by the Benjamini–Hochberg procedure (FDR).

We focused on the Gene Ontology Biological Process category. The results showed strong signal for terms aligned with the input phenotypes, including Leukemia, Atrial septal defect, and Abnormal cardiac septum morphology. The analysis results are shown in Figure 5 and discussed in the main text.

#### 4.4.8 : Method 8 Application: Cross-disease prioritization of genes shared between congenital heart disease and childhood leukemias using graph-based proximity

To identify molecular features shared across congenital and oncologic phenotypes, we applied the Common Neighbors (CN) algorithm from the Neo4j Graph Data Science (GDS) library to measure topological similarity between phenotype and gene nodes in the DDKG.We selected four Human Phenotype Ontology (HPO) terms representing congenital cardiac disorders and pediatric leukemias: Tetralogy of Fallot (HP:0001636), Atrial Septal Defect (HP:0001631), Acute Lymphoblastic Leukemia (HP:0006721), and Acute Myeloid Leukemia (HP:0004808).For each phenotype, we executed the following Cypher query (example shown for ASD):

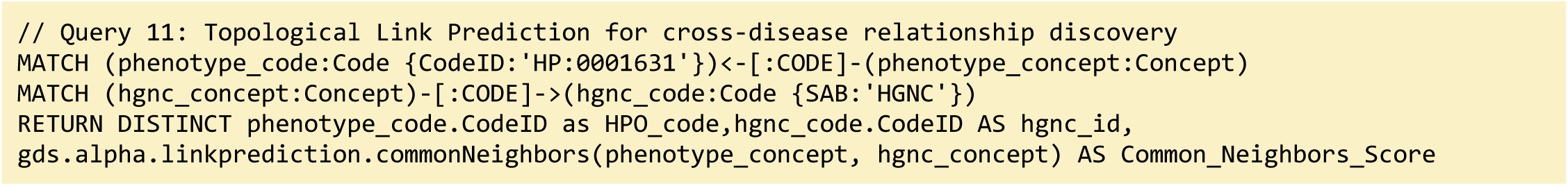

**Query 11 description:** This query computes the number of shared adjacent nodes (Concepts) between each selected phenotype and every human gene Concept in the graph. The query was run independently for all four HPO terms. We utilized the Neo4j Graph Data Science library (gds.alpha.linkprediction.commonNeighbors()) function to generate the Common Neighbors score (CN). The resulting gene–phenotype CN scores were merged into a single results table (see **Table 1**).

Gene–phenotype relationships in the DDKG are derived from integrated sources including OMIM, Orphanet, STRING, and MSigDB, which provide expert-curated gene–disease and pathway annotations. To prioritize genes with potential cross-disease relevance, we applied a two-step rank-based aggregation method. First, for each phenotype, genes were ranked in descending order by CN score (i.e., higher CN score = rank 1).

Second, for each gene, we summed its rank across all four phenotypes. Genes with the lowest total rank were prioritized as top candidates consistently well-connected across both congenital and hematologic disorders.

The top 20 ranked genes are shown in **Table 1** in the main text. These include canonical regulators of both developmental syndromes and cancer pathways, including members of the RAS/MAPK pathway and DNA damage repair systems.

The full list of ranked genes are given in **Supplementary File 3**.

#### Query 11 Semantic Path

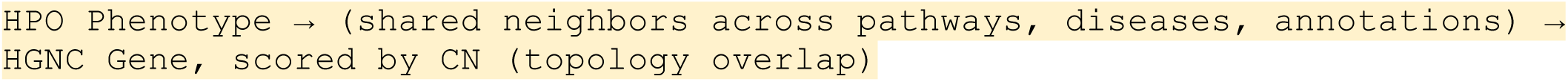

The query defines a semantic comparison between phenotype and gene nodes using the Common Neighbors (CN) algorithm. For each phenotype Concept (HPO-coded), the graph retrieves all human gene Concepts (HGNC-coded) and computes their topological similarity by counting shared adjacent nodes—such as diseases, pathways, interactions, or functional annotations—from integrated resources including OMIM, Orphanet, STRING, and MSigDB.

Even if there is no direct connection between phenotype and gene, the CN score quantifies how densely they are co-embedded in the knowledge graph based on common relationships to other entities. This graph-native computation leverages the semantic richness of the DDKG to identify genes with high contextual overlap with multiple phenotypes. Genes are ranked by their CN scores for each phenotype, and an aggregate rank is used to prioritize candidates most consistently proximal to both congenital and hematologic disease phenotypes.

This path supports cross-disease discovery based on structural integration, not explicit co-annotation, enabling scalable inference of shared gene mechanisms.

##### Validation with external enrichment analysis

To externally validate the DDKG-derived gene rankings, we performed enrichment analysis using the STRING (v12.0) database at https://string-db.org. The top 200 ranked genes were submitted using default parameters and *Homo sapiens* as the reference species. The full submitted list can be found as the first 200 ranked genes in **Supplementary File 3**. STRING applies a one-sided hypergeometric over-representation test to assess whether functional categories occur more frequently in the input gene list than expected by chance, with multiple-testing correction by the Benjamini–Hochberg procedure (FDR).

We focused on the Monarch Human Phenotype Ontology (HPO) category. The results showed strong signal for terms aligned with the input phenotypes, including Leukemia, Atrial septal defect, and Abnormal cardiac septum morphology. The analysis results are shown in Figure 14. These findings support the biological validity of the DDKG-based CN scoring and suggest that topologically prioritized genes represent shared functional mechanisms across congenital and cancer phenotypes. The analysis also provides additional avenues of exploration by ranking other diseases within the gene set.

##### Enrichment analysis within the DDKG

While enrichment analysis was performed externally using STRING for validation for this Method’s case, similar analyses can be implemented directly within the DDKG using graph-native techniques. For example, permutation-based enrichment (sampling) can be conducted by repeatedly sampling random gene sets from the graph while preserving structural properties such as degree distribution or node type. This enables empirical estimation of enrichment baselines against which observed pathway or phenotype annotations can be compared.

Subgraph projection methods could also be applied using Neo4j’s Graph Data Science (GDS) library, where statistical tests could be computed on measured variables including centrality, community membership, or neighborhood density for DDKG Concept classes (e.g., HPO terms, pathways, gene sets). Degree-preserving randomization of phenotype or gene label assignments across the graph structure would allow for testing whether observed overlaps between assertions exceed expectation, such as phenotypes versus gene modules/pathways.

Ongoing work on the DDKG aims to integrate ontology-aware embedding models (using quantities of common neighbor distance or attention-based methods) within the DDKG framework to allow probabilistic inference of pathway or phenotype membership directly from the graph structure itself. These approaches would allow researchers to carry out inference using only the DDKG and its metadata, enabling reproducible, internal validation workflows.

### 4.5 DDKG API: construction and usage

Once the data is ingested, the DDKG provides programmatic access through a safe API, allowing users to interact with the graph without directly accessing the Neo4j database. The API enables secure queries and restricts operations to predefined safe functions, ensuring that users can explore the DDKG’s data in a controlled and reliable manner. This makes the DDKG an ideal platform for both individual researchers and programmatic applications that require access to large-scale biomedical data.

The DDKG API is built on the UBKG API, which uses SmartAPI ^96^. The API offers a structured way to access the Data Distillery Knowledge Graph (DDKG) for programmatic applications. Users of the API can easily integrate multi-omics data into their applications, allowing for cross-dataset analysis, hypothesis generation, and more detailed exploration of molecular relationships. The API is a valuable resource for bioinformatics researchers looking to harness the power of graph databases in a secure and efficient manner.

The API acts as an interface between external applications and the native Neo4j database underlying DDKG, allowing users to retrieve, query, and analyze molecular and biomedical data. However, it is specifically designed to limit the scope of accessible operations, ensuring that users can only perform queries that are safe and predefined by the DDKG API. This makes it a secure and controlled access point, reducing the risk of unintended queries on the DDKG. Information on the UBKG API can be found at https://ubkg.docs.xconsortia.org/api/.

The DDKG API’s design enables safe integration into larger programmatic workflows. It allows researchers and developers to build applications that use the graph for advanced data mining, link discovery, and machine learning tasks without needing direct access to the full capabilities of the underlying Neo4j database. This safety-focused design ensures that all queries and operations remain within the boundaries defined by the DDKG API, providing a secure environment for computational analysis while maintaining the integrity of the knowledge graph.

### 4.6 Visualization Portal: Development of the DDKG-UI

The Data Distillery Knowledge Graph user interface (DDKG-UI) was developed with Next.JS, React, and Material UI to provide an open source web-based interface to the DDKG. To visualize subnetworks returned from Neo4j from Cypher queries, ball-and-stick subnetworks are visualized with Cytoscape.JS ^97^. The DDKG-UI hosts a subset of the entire DDKG. It serves assertions from the 11 Common Fund programs as well assertions from ClinVar ^98^, HGNC ^89^, HPO ^99^, UniProt ^100^, and MSigDB ^101^. Assertions based on gene-gene co-expression correlations from ARCHS4 ^102^ are included to draw direct connections between genes (**Table 4**). The DDKG-UI leverages the same framework developed for ReproTox-KG ^8^, Enrichr-KG ^103^, and Harmonizome-KG ^104^.

**Table 4.**
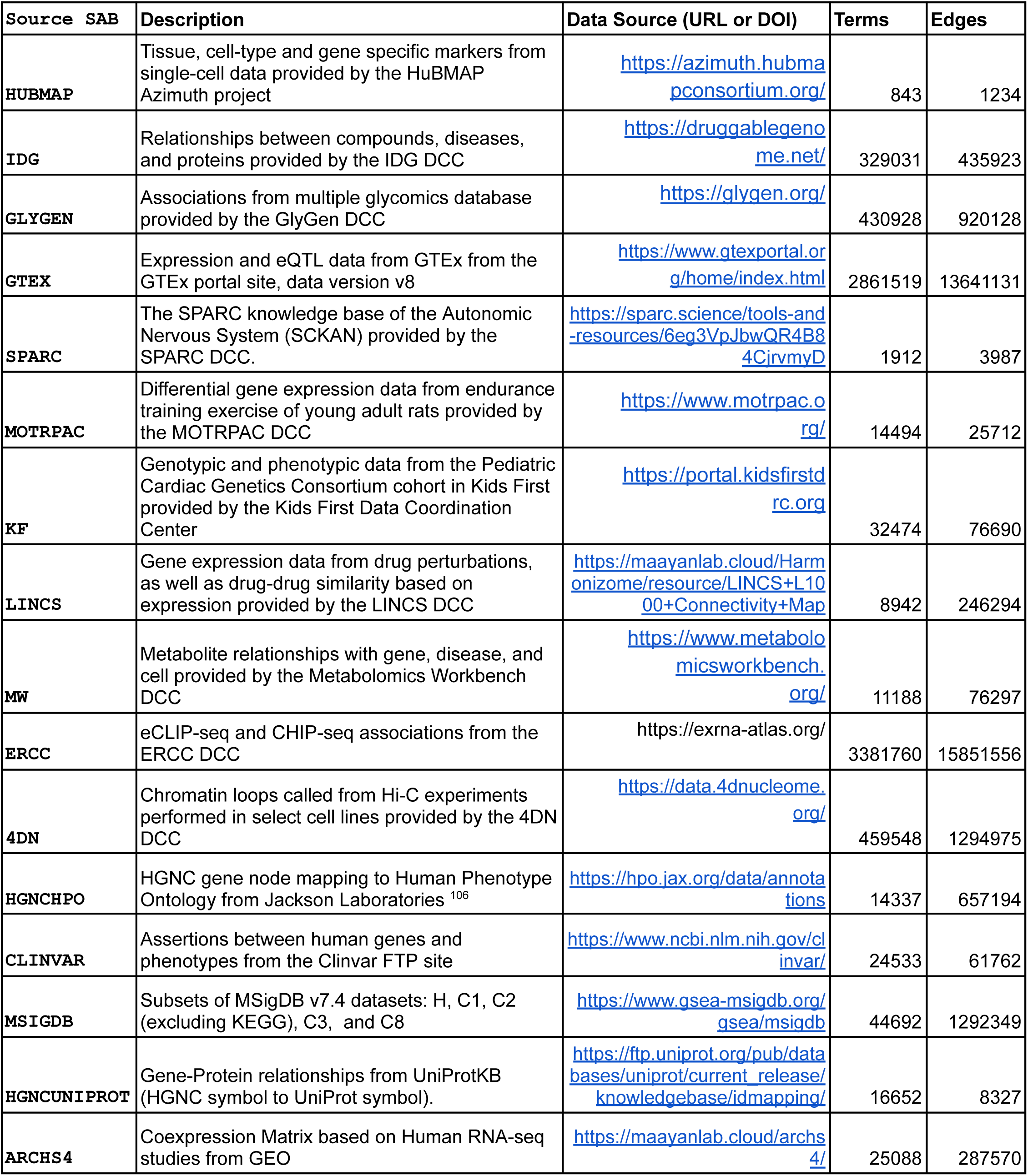
Assertions from the DDKG specifically available within the subgraph exported to DDKG-UI. DCC=Data Coordination Center.

To adapt the UBKG serialization to be compatible with DDKG-UI, Concept, Code, and Term nodes were collapsed into a singular node with additional metadata. For each Concept node, we took the connected Term node and its label as the node’s label. We then added the associated ontology and controlled vocabulary identifiers from the linked Code nodes. This information was added as the new node’s metadata. To ensure consistency across identifiers, we converted the `Concept` identifiers to uuids using UUID5. Lastly, node types are manually determined based on their source. Since the assertions were contributed by the different Common und DCCs, this information is obtained from the DDKG’s Data Dictionary. We illustrate a simple example of this conversion with the drug, Dexamethasone (Figure 15). The nodes are then saved into a CSV file that lists them. For the edges, the triples from the CUI-CUI.tsv file, which are links between Concepts, are converted into an edge list that is also saved as a separate CSV file. For this, we filtered the file to include only edges that are from selected sources (**Table 4**).

DDKG-UI also includes assertions between genes and their functions predicted based on gene-gene co-expression correlations from RNA-seq datasets ^102^. First, we generated gene-gene co-expression edges by taking the top ten co-expressed genes for each human gene in DDKG. Next, we took the different non-gene terms and linked them to their respective gene sets in Enrichr ^105^. By computing the mean correlation coefficient between the associated gene set top 10 most co-expressed genes we created a new set of edges. These edges are added to the DDKG-UI as predicted links. These additional links were converted into assertions and were ingested into the Neo4j DDKG-UI instance. The DDKG-UI provides means to search the DDKG by term search with autocomplete, as well as by two-term search to construct subnetworks that connect them. In addition, five use cases have dedicated pages that invoke the Cypher query dedicated to these use cases. The DDKG-UI also hosts a web-based version of the Data Dictionary as well as download, tutorial, and API documentation to enable users to utilize the resources for other applications.

## Acknowledgements

Support from the NIH Common Fund through the Office of Strategic Coordination/Office of the NIH Director under awards R03OD030600, OT2OD030162, and U24OD038422 to D.M.T., B.S., T.M.A., D.D.H., J.T., C.N., and A.R., with additional institutional funding from the Children’s Hospital of Philadelphia Research Institute.

J.C.S. and J.A.S. acknowledge support from the NIH Common Fund through the Human BioMolecular Atlas Program (HuBMAP, OT2OD033759), the Common Fund Data Ecosystem Supplement (CFDE, OT2OD030545), and the SenNet program (U24CA268108).

J.J.Y., C.G.L., and V.T.M. were supported by the Illuminating the Druggable Genome (IDG) program under NIH award OT2OD030546.

A.M., J.E.E., D.J.B.C., Z.X., H.K., and S.L.J. were supported by the LINCS and Data Resource Center initiatives under awards U24CA271114, U24CA264250, OT2OD036435, and OT2OD030160.

S.S., M.R.M., S.R., and E.F. acknowledge support from the Metabolomics Workbench (MW) and related programs under awards OT2OD030544, OT2OD036435, U2C-DK119886, and U24DK141185.

Support for the SPARC program was provided to B.d.B., J.S.G., T.H.G., F.T.I., and N.K. under awards OT2OD030541 and OT2OD032619.

M.E.R., R.F., D.J., A.Mi., and A.Mil. acknowledge support from the ERCC and related efforts under awards OT2OD030547, U54DA049098, and U01AG072439.

R.M., J.V., M.T., and R.R. were supported by the GlyGen and BiomarkerKB initiatives under NIH awards OT2OD032092, 1R24GM146616-01, and U24OD038423.

P.P., A.J.S., C.B., and J.M. acknowledge support from the 4D Nucleome (4DN) program under NIH award U01CA200059.

